# A unified intermediate-resolution framework for modeling biomolecular condensates at near-cellular complexity

**DOI:** 10.64898/2026.09.15.751913

**Authors:** Shanlong Li, Jessica Fong Ng, Jianhan Chen

## Abstract

Biomolecular condensates, formed through phase separation of multivalent biomolecules, are central to cellular organization and function. Here, we present a unified, physics-based intermediate-resolution framework for efficient and quantitative simulation of biomolecular phase separation at near-cellular complexity. The framework integrates the HyRes protein and iConNA nucleic acid models to explicitly describe major driving forces and capture transient local and global structural features underlying phase transitions. With well-balanced hydrogen bonding, electrostatic, cation-π, and hydrophobic interactions, as well as nucleic acid base stacking, base pairing, and ion-mediated effects, the framework accurately predicts saturation concentrations of over 60 disordered proteins (within tenfold) and phase diagrams of homotypic RNA and heterotypic protein-nucleic acid condensates. We further introduce iConMetabolome, a machine learning-enabled library of cellular metabolites that captures metabolite-driven phase separation and reproduces experimental partitioning trends across condensates. This unified platform enables the dissection of the mechanisms and regulation of condensation in biology and disease.

## Introduction

The recognition that phase separation is the defining mechanism for the formation of cellular membraneless organelles has generated intense interest in recent years^1–3^. Phase separation in the cell is mainly driven by dynamic and multivalent macromolecules including intrinsically disordered proteins (IDPs), flexible RNAs, and DNAs^4–6^. It gives rise to compositionally distinct bodies, referred to as biomolecular condensates^7^. They have been shown to contribute to a myriad of biological functions including RNA processing, DNA repair, stress responses, trafficking, metabolism, and cellular signaling^8–11^. Within the cell, condensates are remarkably complex multicomponent systems and could consist of many distinct proteins, nucleic acids, and small molecules, particularly metabolites^8,12^. Different client macromolecules and small molecules can selectively partition into or be excluded from specific condensates, establishing distinct microenvironments to support function^13–15^. Moreover, condensates are dynamically regulated and can undergo aging and maturation, transitioning from liquid-like to gel or solid-like states with potentially pathological consequences in neurodegenerative diseases^16,17^. Despite significant advances, much remains to be understood about the molecular driving forces and regulation of condensation and how condensate properties are modulated to support specific cellular functions or become dysregulated in disease.

Molecular simulation is indispensable for complementing experiments in mechanistic studies of biomolecular condensates^18,19^. To access the large length (> ~100 nm) and time scales (> μs) required for direct simulation of phase transitions, many coarse-grained (CG) models at one-bead-per-residue resolution have been developed for proteins, including the hydropathy scale (HPS) models^20,21^, CALVADOS^22^, Mpipi^23^, MOFF^24^, and COCOMO^25^. These models have enabled important advances in dissecting sequence-dependent phase behaviors of IDPs. Compatible nucleic acid models have been introduced at one or two beads per nucleotide, such as HPS-RNA^26^, COCOMO^25^, and CALVADOS-RNA^27^, allowing initial investigations of protein-nucleic acid co-phase separation. Independent CG models with 1-3 beads have also been developed for RNA phase separation, such as SIS^28^, RNA2PS^29^, and others^30^. Multi-bead CG protein models, including the popular MARTINI force field, have also been explored for simulation of IDP phase separation^31–33^. A fundamental limitation of these models is that low resolution inherently precludes the description of protein backbone hydrogen bonding and transient secondary structures that are prevalent in IDPs and important in condensation^34^. Similarly, one- or two-bead nucleic acid representations cannot simultaneously capture diverse RNA interactions including phosphate electrostatics, base stacking, base pairing, and divalent ion-mediated effects that are central to RNA condensation and its regulation. These fundamental limitations point to the need for higher-resolution CG models that balance computational efficiency with sufficient chemical detail to describe the rich balance of distinct interactions and transient structures of both proteins and nucleic acids in condensation.

To address these limitations, we have developed a hybrid resolution (HyRes) protein model for reliable description of local and long-range structures of IDPs^35–37^. The backbone is represented at the all-atom level to capture backbone-mediated interactions and secondary structures, and side chains are represented at intermediate resolution with up to 5 beads, allowing semi-quantitative description of transient long-range organizations (**Figure 1a**). With the latest optimization, HyRes accurately predicts residual helicity profiles of an independent test set of 40 IDPs with ~0.83 correlation, captures the radii of gyration (*R*g) for a large test set of over 100 IDPs with ~0.97 correlation, and recapitulates NMR paramagnetic relaxation enhancement (PRE) profiles of nine IDPs at a level comparable to or better than the latest all-atom force fields^37^. Initial applications to IDP phase separation further demonstrated that HyRes could capture the coupling between secondary structure and phase separation of IDPs, correctly predicting the effects of single mutations on TDP-43 helicity and phase separation propensities^38^. We further designed intermediate-resolution models for condensates of RNAs (iConRNA)^39^ and DNAs (iConDNA)^40^, collectively referred to as iConNA herein, that can capture local and long-range structural features of dynamic RNAs and DNAs and their complex phase behaviors. Representing each nucleotide with 6-7 beads (**Figure 1b**), iConNA explicitly describes distinct interactions of the phosphate backbone and nucleobases, such that one can faithfully capture the conformational dynamics of nucleic acids and study the detailed interplay of base pairing, base stacking, electrostatics, ion interactions, and transient local structures in phase separation. iConRNA, in particular, has been shown to recapitulate the lower critical solution temperature (LCST) phase separation of RNAs and its nontrivial dependence on sequence, length, ion and RNA concentrations, and Mg^2+^ level^39^. This is highly noteworthy and has not been achieved by any other atomistic or CG models of RNAs.

**Fig. 1.**
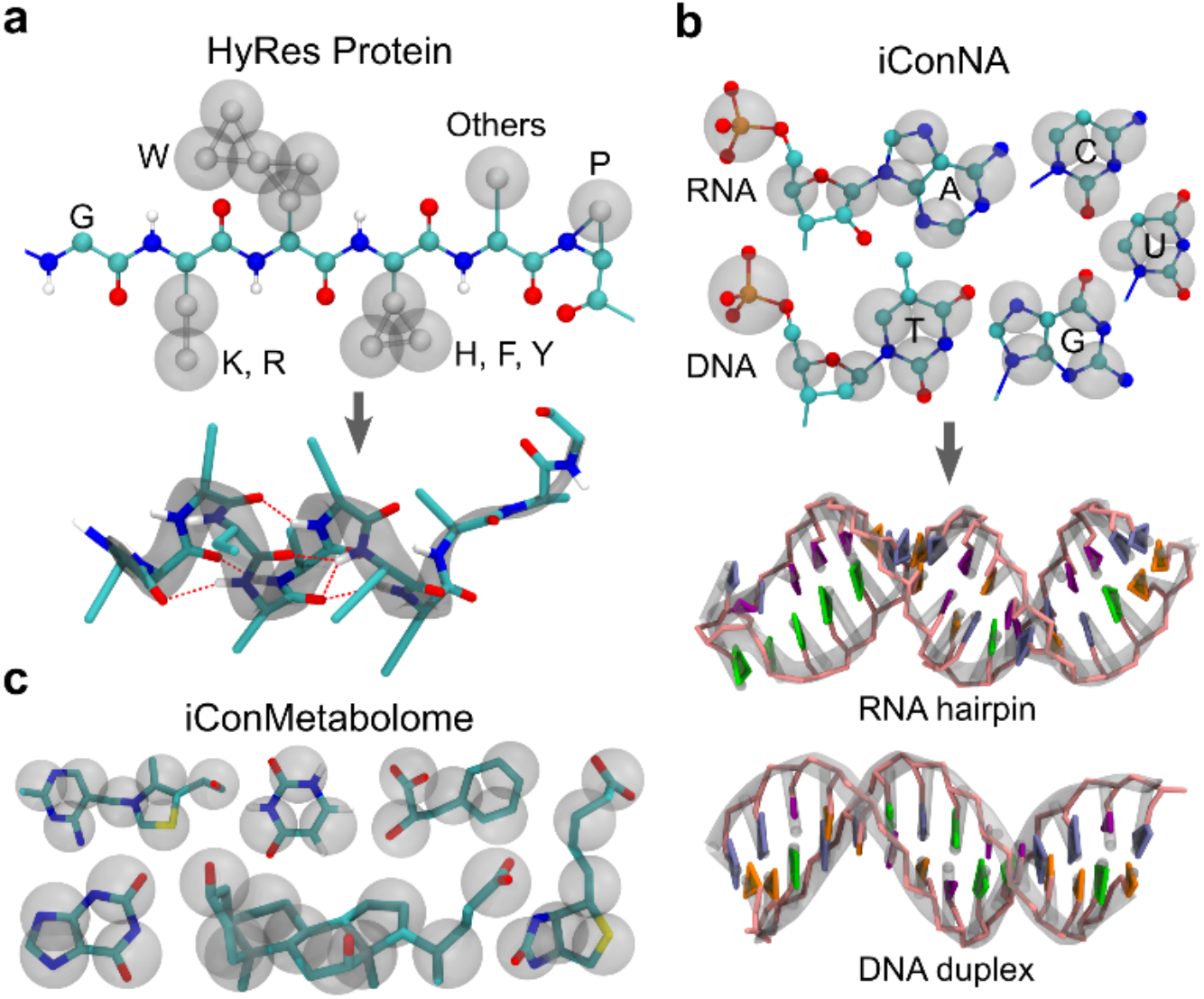
Representations of HyRes, iConNA, and iConMetabolome. **a-c**, CG representations and representative structures of (**a**) HyRes for proteins, (**b**) iConNA for RNAs and DNAs, and (**c**) iConMetabolome for cellular metabolites. The atomistic backbone of HyRes enables accurate modeling of backbone-mediated hydrogen bonding and transient secondary structures, while the intermediate resolution of iConNA allows explicit treatment of diverse backbone charge interactions, base stacking, and base pairing to capture dynamic structural and thermodynamic properties of nucleic acids.

In this work, we integrate HyRes and iConNA to derive a unified, physics-based framework for efficient and accurate simulation of homotypic and heterotypic protein and nucleic acid phase separation. Through extensive experimental data-guided optimization, we show that the HyRes/iCon framework achieves accurate prediction of the conformational properties and phase behaviors of diverse intrinsically disordered and multidomain proteins. It also captures the dimensions, folding, and condensation of structured and flexible nucleic acids. Critically, the integration enables quantitative simulation of heterotypic phase transitions, reproducing experimental phase diagrams and material properties across a range of protein-RNA and protein-DNA systems. We further extend the framework to the broader cellular environment by introducing iConMetabolome, a machine learning (ML)-enabled library of small-molecule models that captures both metabolite-driven phase transitions and metabolite partitioning properties in condensates. Together, these advances establish the integrated HyRes/iCon framework as an efficient platform for simulating biomolecular condensation at near-cellular complexity with quantitative accuracy.

## Results

### Quantitative simulation of IDP phase separation

The atomistic backbone in HyRes enables accurate modeling of anisotropic hydrogen bonding interactions and transient secondary structures, while the intermediate resolution side chains support efficient description of diverse physical interactions and transient long-range contacts (**Fig. 1a**). Benchmarks against over 100 IDPs demonstrated that the latest HyRes generates atomistic ensembles of the monomers similar to and often better than those from state-of-the-art all-atom protein force fields in reproducing experimental chain dimensions, secondary structures, and transient tertiary contacts, but at several orders of magnitude lower computational cost^37^. To further evaluate the ability of HyRes to simulate protein phase separation, we compiled a large set of 64 experimentally measured saturation concentrations (*C*_sat_) for 13 wild-type (WT) IDPs and their variants (**Supplementary Table S1**), and performed large-scale phase co-existence simulations in HyRes to determine *C*_sat_ (see **Methods**). These IDPs include short model peptides to complex cellular proteins, spanning a broad range of sequence lengths (*L* = 10-284 aa) and diverse amino acid compositions (**Extended Data Fig. 1a**). They also exhibit a wide range of charge-patterning characteristics, with κ values ranging from 0 (well-mixed, *e*.*g*., GY-23) to 1.0 (fully segregated, *e*.*g*., K10/D10). The experimental *C*_sat_ span almost 4 orders of magnitude, from low µM to several mM, measured at temperatures ranging from 277 to 298 K. Collectively, this diverse benchmark set provides a stringent test of the balance among van der Waals, polar-polar, π-π, cation-π, and electrostatic interactions encoded in the HyRes model.

As summarized in **Fig. 2a**, calculated *C*_sat_ values mostly fall within 10-fold of the experimental results (shaded region), with an overall Pearson’s correlation of *r* = 0.76. When considering the WT sequences only, the correlation is much higher at *r* = 0.93 (**Fig. 2b**). To the best of our knowledge, this level of correlation across such a large and diverse *C*_sat_ benchmark has not been demonstrated by any other CG or atomistic approaches; it is especially noteworthy considering the sequence diversity and large dynamic range of *C*_sat_. The ability of HyRes to qualitatively predict the mutational effect on phase separation propensity is also noteworthy, as illustrated using the set of 24 A1-LCD variants (**Extended Data Fig. 1b**). We further illustrate the efficiency of HyRes and the level of convergence achieved using the 151-residue TDP-43 LCD as a representative system. With 100 copies of TDP-43 LCD in a 100-nm cubic box (~90K beads, ~170 µM), the system reached stable phase co-existence during the 4 µs simulation, which took only ~2 days on a single Nvidia L40S GPU. Analysis showed that essentially all structural properties seem fully converged within the condensate, with all protein chains exhibiting nearly identical *R*_g_ and end-to-end distance (*R*_ee_) distributions (**Extended Data Fig. 1c**) as well as residue helicity profiles (**Extended Data Fig. 1d**). The intermolecular contact probability calculated from the first and second µs segments of the last 2 µs trajectory show only minor differences, with relative deviations below ~10% (**Extended Data Fig. 1e**).

**Fig. 2.**
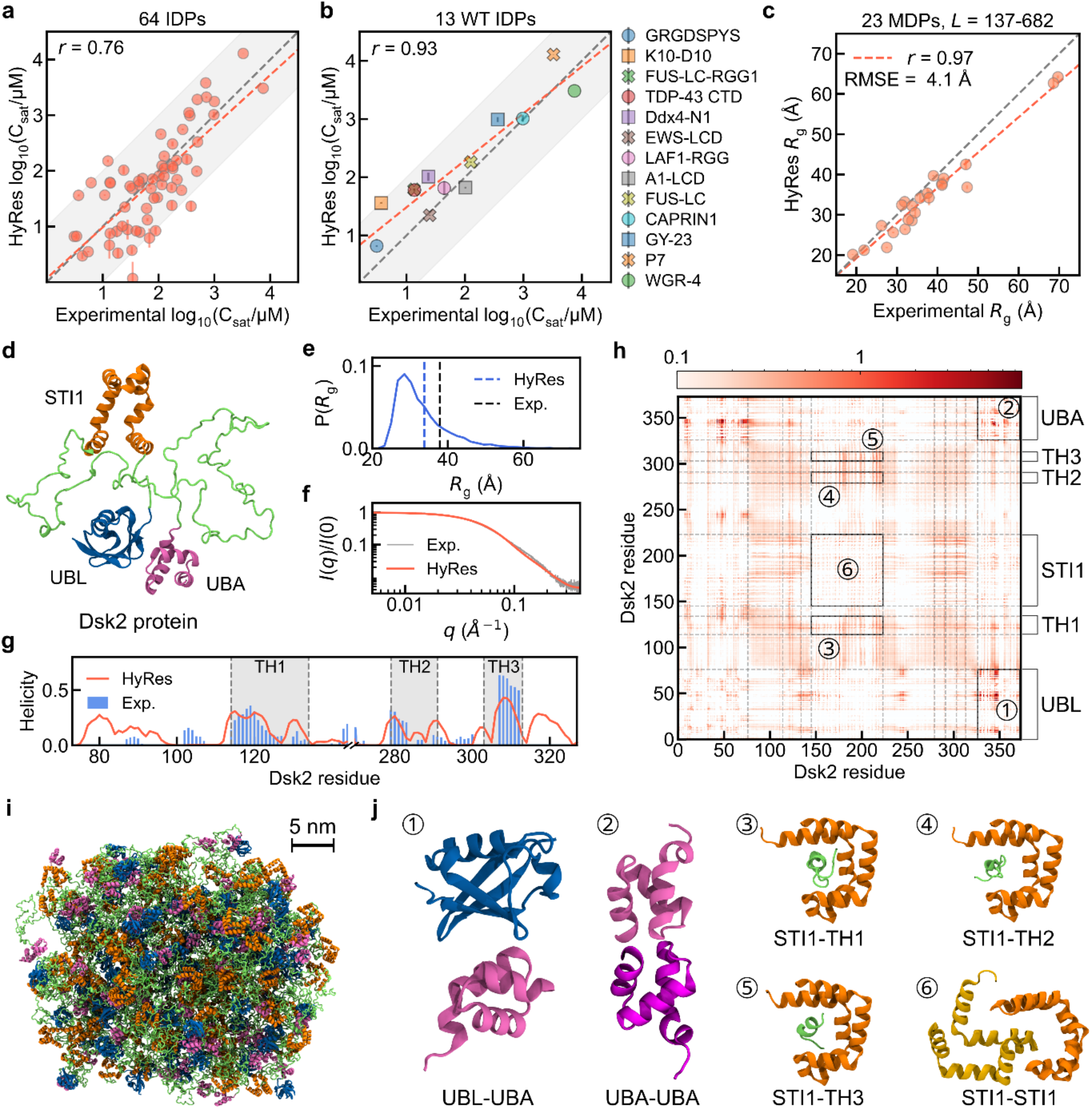
Accurate simulation of protein phase equilibrium using HyRes. **a,b**, Comparison of HyRes-derived and experimental *C*_sat_ for 64 IDPs (**a**) and 13 WT IDPs (**b**). The gray shaded region indicates values within 10-fold of the experimental values. Grey dashed diagonal lines plot *y* = *x*, and red dashed lines plot the best linear fit. Error bars in the calculated values were estimated from block analysis. **c**, Comparison of HyRes-derived and experimental *R*_g_ for 23 MDPs. **d**, A representative snapshot of Dsk2, showing its folded domains, UBL (blue), UBA (mauve), and STI1 (orange), and disordered segments (lime). **e,f**, Comparison of HyRes-derived and experimental dimensions (*R*_g_, **e**) and scattering profiles (**f**) of protein Dsk2. **g**, Comparison of HyRes- and NMR-derived residual helicities of the disordered segments of Dsk2. Transient helical domains 1, 2, and 3 (TH1, TH2, and TH3) are highlighted in grey. **h**, Intermolecular residue-residue contact map derived from the condensate simulation (see Methods). For each residue pair, the contact density represents the average number of contacts within condensates. Contacts with density below 0.1 were masked (white). Rectangular areas 1-6 and the corresponding representative snapshots (**j**) highlight the interactions contributing to Dsk2 phase separation. **i**, A representative snapshot of a condensate of 98 Dsk2 molecules, using the same coloring scheme as in **d**.

### Accurate simulation of multi-domain protein conformation and condensates

We next evaluated the ability of HyRes to describe dynamic interactions of both folded and disordered domains. For this, we collected 23 multidomain proteins (MDPs) with available experimental *R*_g_ values that range from 137 to 682 residues in length^41,42^ (**Supplementary Table S2**). With the folded domains treated as rigid bodies, HyRes simulations predicted *R*_g_ with a strong correlation of *r* = 0.97 and a small root mean squared error (RMSE) of ~4.1 Å compared to SAXS-derived values (**Fig. 2c** and **Extended Data Fig. 1f**). This suggests that the interactions between folded and disordered domains are well balanced in HyRes. We then applied HyRes to examine the structural dynamics and phase separation of the yeast protein Dsk2, a ubiquitin-binding shuttle protein that scaffolds proteasome-containing condensates under stress conditions^43^. The 373-residue protein contains three folded domains, namely, the ubiquitin-like (UBL, residues 1-76), ubiquitin-associated (UBA, residues 326-370), and stress-inducible 1 (STI1, residues 145-223) domains, in addition to disordered regions (**Fig. 2d**). HyRes accurately captures the overall dimension of Dsk2 and its scattering profile (**Fig. 2e-f**), with a modest underestimation of *R*_g_ (33.1 versus 37.9 Å), and all three transient helical (TH) domains identified by NMR (**Fig. 2g**)^44^.

Several distinct multivalent interactions have been suggested to drive Dsk2 phase separation, including UBL-UBA, STI1-STI1, and STI1-TH interactions^44^. Simulation of a droplet containing 98 Dsk2 molecules recapitulated the importance of these interactions and revealed additional multivalent contacts (**Fig. 2h-j)**. The specific UBL-UBA interaction (➀) emerged as a prominent interaction, whereas the UBA-UBA interaction (➁) was also frequently present in the condensate^45^. Representative snapshots showed diverse and dynamic UBL-UBA and UBA-UBA binding configurations (**Supplementary Fig. S1a-b**). The interactions between the STI1 domain and each of the three TH domains were also captured (➂‐➄), with TH1-3 transiently engaging the STI1 pocket through both helical and disordered conformations (**Supplementary Fig. S1d**), which is highly consistent with experiment^44^. In addition, STI1-STI1 self-association was observed in multiple configurations (➅; **Fig. 2j** and **Supplementary Fig. S1c**), although these interactions were weaker and more dynamic. Beyond these specific interactions, the two disordered regions (residues 77-144 and 224-325) formed extensive interactions with themselves, with each other, and with STI1 domains, which likely provide an additional major contribution to Dsk2 phase separation. The ability of HyRes to accurately simulate MDP phase separation is further supported by co-phase separation of polySH3 and polyPRM proteins, which will be discussed in a later section.

### iConNA accurately samples flexible nucleic acids and their phase transitions

Representing each nucleotide with six or seven CG beads (**Fig. 1b**), iConNA is able to capture the complex interplay of distinct backbone and side chain interactions, including base stacking, base pairing, screened electrostatics, and explicit Ca^2+^ and Mg^2+^ ions^39,40^. The original iConRNA model was parameterized with a dielectric constant of 20. To integrate with HyRes, we first increased the dielectric constant in iConRNA to 60 (at room temperature), followed by systematic calibration to rebalance various interactions as done previously^39^. The refined iConRNA model accurately reproduces the dimensional properties of homopolymer RNAs, including the *R*_g_ of rA_30_ and rU_30_ and persistence length (*L*_p_) of rU_40_, even though *R*_ee_ remains over-estimated for rU_40_ (**Fig. 3a**). The calculated scattering profiles of both rA_30_ and rU_30_ agree well with experimental data (**Extended Data Fig. 2c**). The refined model also improved the description of local and global RNA conformational properties relative to the original iConRNA model (**Fig. 3a**; **Extended Data Fig. 2a-b**). We further benchmarked the thermodynamic stabilities of RNA secondary structures using eight hairpin variants at 150mM NaCl and three duplexes at 1, 5, 10, and 50 mM MgCl_2_ with 25 mM NaCl (**Fig. 3b-c** and **Supplementary Table S3**). The predicted melting temperatures showed strong agreement with experimental results, with an RMSE of 3.5 °C (*r* = 0.95) for the hairpins and 0.6 °C (*r* = 0.96) for the duplexes, respectively, suggesting that both base pairing and stacking interactions are well balanced.

**Fig. 3.**
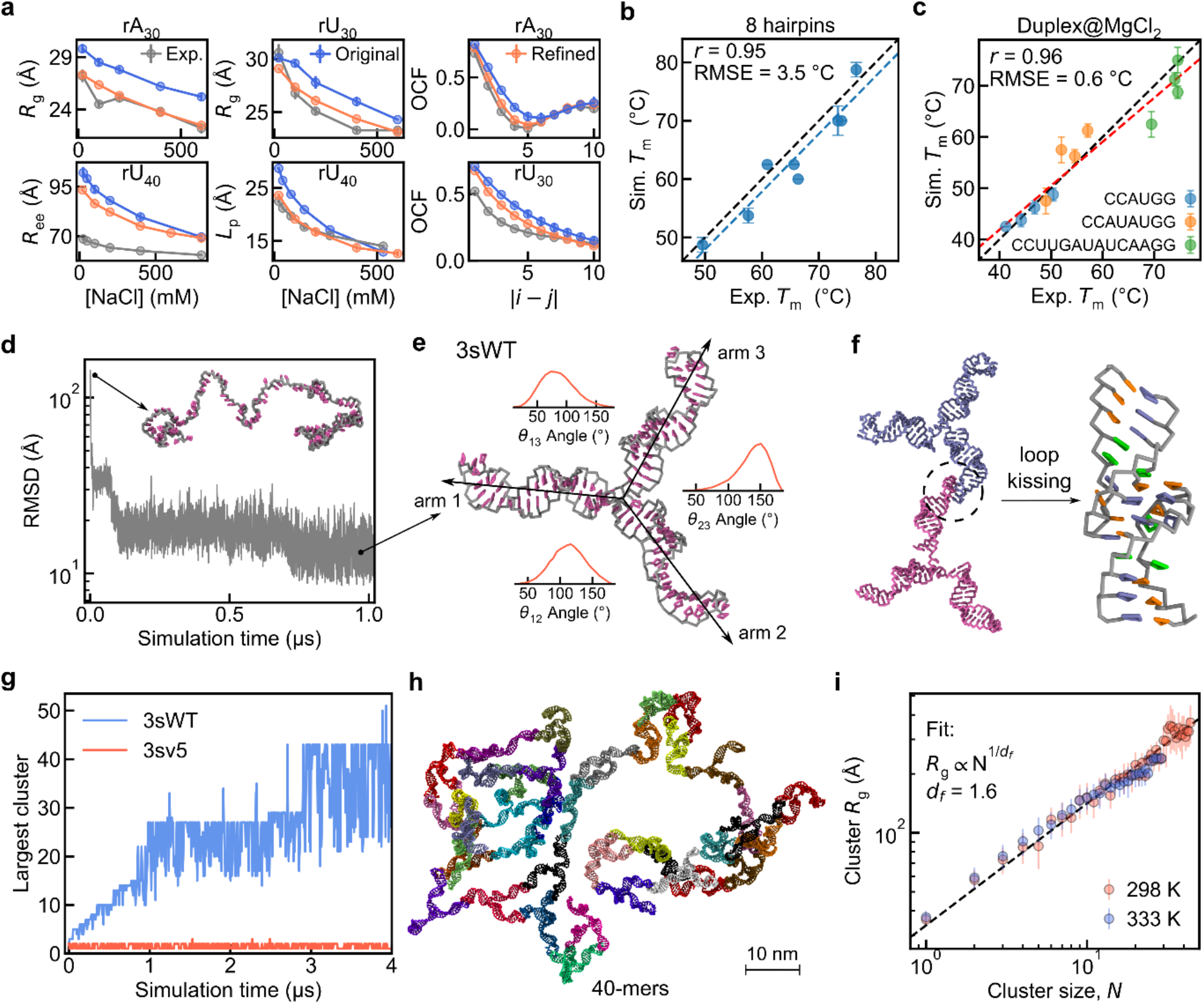
Revised iConRNA and simulations of RNA nanostar phase separation. **a**, Comparison of homopolymer RNA dimensional properties from experiments, original and refined iConRNA models. Error bars were estimated from block analysis. **b,c**, Correlation between calculated and experimental melting temperatures (*T*_m_) of (**b**) 8 RNA hairpin variants and (**c**) three RNA duplexes under four MgCl_2_ concentrations. In **b** and **c**, error bars were estimated from two independent replicates. **d**, Folding trajectory of the 3sWT nanostar from a random-coil conformation (shown as the insert). Only the first half of the 2-µs trajectory is shown for clarity. **e**, A representative folded structure of the 3sWT nanostar, with the distributions of the angles between pairs of arms also shown. **f**, A representative kissing-loop interaction between two 3sWT nanostars. **g**, Largest cluster sizes as a function of time during 4 µs phase separation simulations of 3sWT nanostar vs the non-phase-separating 3sv5 variant. **h**, A representative network of 40 3sWT nanostars. **i**, Fractal scaling of cluster size (in *R*_g_) to the number of RNA molecules (*N*) in the cluster. A consistent scaling relationship was observed across temperatures. The dashed line indicates the fitted relationship, *R*_g_ ∝ *N*^1/*d*f^, where *d*_f_ is the fractal dimension (~ 1.6 here). Error bars indicate the range of *R*_g_ among clusters containing *N* nanostars.

iConNA can recapitulate the complex dependence of nucleic acid phase separation on sequence, length, concentration, temperature, and Mg^2+^ level^39^. Here we further evaluate the ability of iConRNA to fold 3-arm RNA nanostars and capture their phase separation driven by loop-loop interactions^46^. Specifically, we focus on the single-stranded 3sWT RNA that contains the human immunodeficiency virus (HIV) kissing loop (KL) (GCGCGC), as well as 5 KL variants (3sv1-5, **Supplementary Table S4**). Starting from fully extended conformations, all six sequences readily folded into stable 3-arm nanostar structures (**Fig. 3d-e** and **Extended Data Fig. 2e**). Analysis of the angles between pairs of arms revealed significant flexibility and intriguing deviations from the ideal threefold symmetry. While the star structure is mostly planar, the angle between arms 2 and 3 is significantly larger (~150°) than the other two (~90° between arms 1 and 3) (**Fig. 3e** and **Extended Data Fig. 2f**). With 100 copies in a 120-nm cubic box (~100 μM), 3sWT RNAs readily form fully paired KL interactions (**Fig. 3f**), which phase separate to into an extended and dynamic RNA cluster of up to ~40 molecules (**Fig. 3g-h**). In contrast, the 3sv5 variant with GCUCGC loops (and thus weakened KL interactions) remained dispersed throughout the 4-µs simulations (**Fig. 3g**), fully consistent with experiments^46^. We further characterized the architecture of the 3sWT condensates by examining the fractal scaling relationship between the cluster dimension in *R*_g_ and the number of RNA molecules (*N*), *R*_g_ ∝ *N*^1/*d*f^, where *d*_f_ is the fractal dimension^47^. At both 298 and 333 K, the clusters exhibited a consistent fractal dimension of *d*_f_ = 1.6 (**Fig. 3i**), identical to the effective fractal dimension reported for DNA nanostar networks measured by SANS^48^. This value differs markedly from those of rA_30_ (*d*_f_ = 3.1) and the CAPRIN1 protein (*d*_f_ = 3.6) (**Extended Data Fig. 2g**), indicating the distinct network architecture of nanostar condensates. Notably, *d*_f_ = 1.6 is also consistent with the theoretical prediction for the percolation of diluted, swollen branched polymers^49^, suggesting a potential coupling between phase separation and percolation in the formation of RNA nanostar condensates. That iConRNA accurately captures the structure, fractal dimension, and stability of RNA nanostar condensates further supports the balanced representation of competing interactions in the model.

### HyRes/iConNA for heterotypic phase separation of protein and nucleic acids

Besides recalibration of iConRNA using a dielectric constant of 60, integration of HyRes and iConNA requires balancing interactions between amino acid side chains and nucleic acid bases. Taking Arg-Arg interaction as the baseline, pairwise interaction potentials were determined according to the relative strengths derived from recently reported all-atom calculations^50^ (see **Methods** and **Supplementary Table S6**), followed by validations against several protein-nucleic acid condensation systems. The balance of protein-protein, protein-nucleic acid, and nucleic acid-nucleic acid interactions was first examined by studying the phase behavior of poly(Pro-Arg)_30_ ((PR)_30_) peptides and homopolymer RNAs (rA_30_ and rU_30_), where the phase diagrams as a function of concentration and ionic strength have been determined experimentally^51^. The results show direct phase coexistence simulations using HyRes/iConNA precisely reproduced the experimental phase boundary for rA_30_, even though the salt concentrations required to dissolve rU_30_/(PR)_30_ condensates were slightly over-estimated (**Fig. 4a, Extended Data Fig. 3a**). Contrast-variation solution X-ray scattering (CV-SAXS) measurements showed that both rA_30_ and rU_30_ become more flexible in the presence of (PR)_30_, as indicated by a decrease in the Flory’s scaling exponent (*ν*)^51^. The same trend was reproduced by the simulations (**Fig. 4b-c**), which can be attributed to electrostatic screening of the negatively charged RNA backbone by positively charged peptides. Interestingly, the simulations also reproduced the curious increase in *R*_g_ accompanying the decrease in *v* observed experimentally. Analysis of RNA conformations revealed that the apparent chain expansion arises from larger *L*_p_ (**Extended Data Fig. 3b**), which can be quantified by fitting the chain conformations to the helical worm-like chain (HWLC) model^52^. In the presence of (PR)_30_, screening of electrostatic self-repulsion along the RNA backbone increases backbone flexibility, resulting in a lower *v*. At the same time, base-base interactions are enhanced, increasing the average neighboring phosphate distance (*L*_b_) (**Extended Data Fig. 3b**) and consequently *L*_*p*_. Following 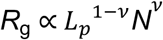, the concomitant decrease in *v* and increase in *R*_g_ could be rationalized by the increase in *L*_p_.

**Fig. 4.**
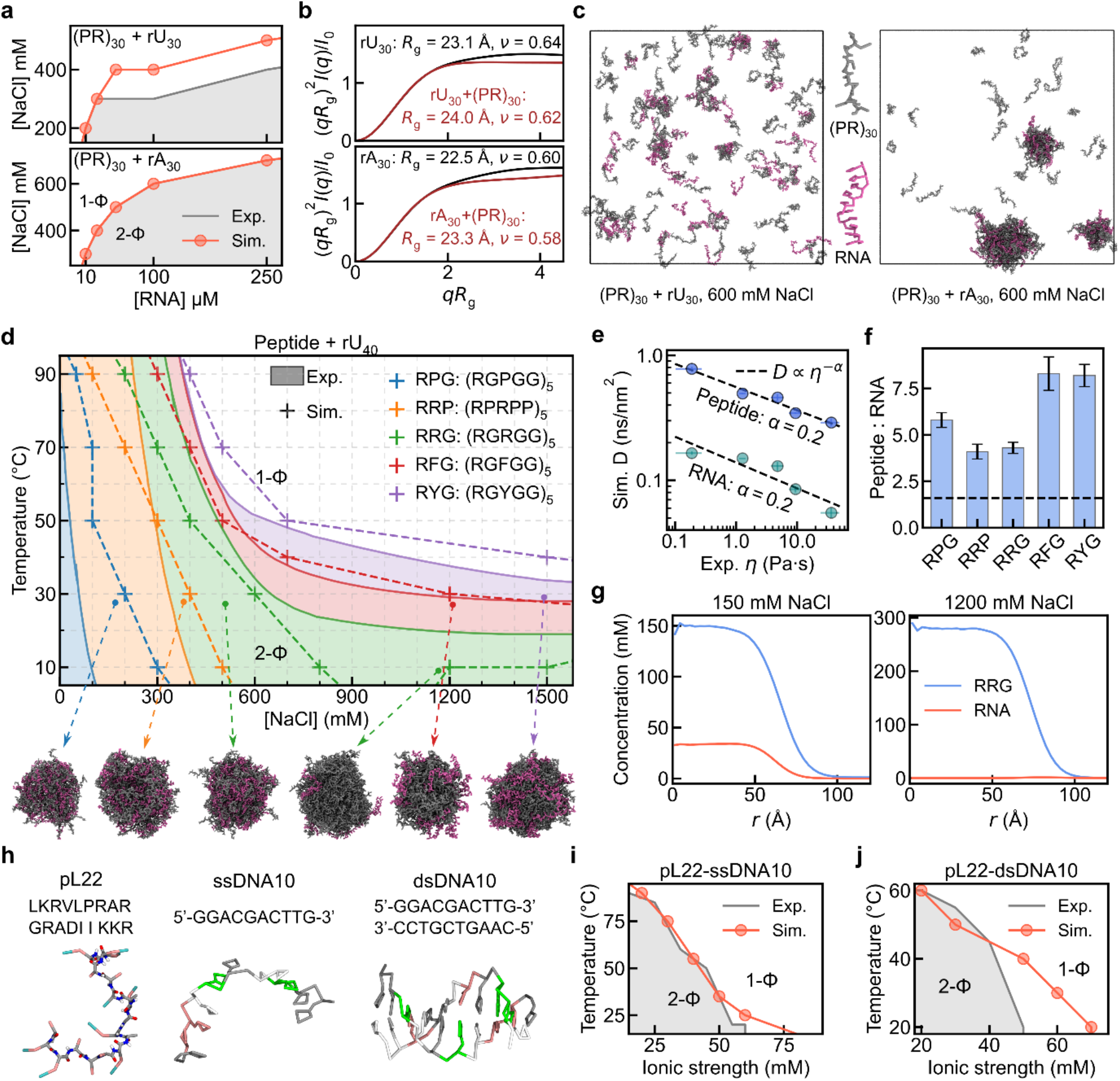
HyRes/iConNA for heterotypic phase separation of proteins and nucleic acids. **a**, Comparison of simulated and experimental phase diagrams for (PR)_30_-rU_30_ and (PR)_30_-rA_30_ mixtures. The peptide: RNA stoichiometric ratio was fixed at 2:1. **b**, Simulation-derived scattering profiles reproduce the experimentally measured changes in RNA dimensions within condensates at 600 mM NaCl. **c**, Representative snapshots of (PR)_30_-rU_30_ and (PR)_30_-rA_30_ mixtures at 600 mM NaCl. **d**, Comparison of simulated and experimental phase diagrams of rU_40_ with model peptides RPG, RRP, RRG, RFG, and RYG as a function of temperature and salt. The peptide and RNA concentrations are 5.0 and 2.5 mg/ml. Representative condensates observed under the conditions indicated by arrows are shown below. Peptide and RNA chains are shown in grey and mauve, respectively. **e**, Correlation between the simulated diffusion coefficient (*D*) and experimental viscosity (*η*). Dashed lines indicate linear fits of log(*D*) versus log(*η*), with correlations of *r* ~ −0.98 and −0.91. **f**, Peptide-to-RNA ratio within condensates. The dashed line indicates the stoichiometric ratio for charge neutralization. **g**, Density profiles of peptide and RNA molecules for RRG-RNA condensates at 150 and 1200 mM NaCl. **h**, Sequences and representative snapshots of pL22, ssDNA10, and dsDNA10. **i, j**, Comparison of simulation-derived and experimental phase diagrams for pL22-ssDNA10 (**i**) and pL22-dsDNA22 (**j**) mixtures. The concentrations of pL22, ssDNA10, and dsDNA22 are 300, 220, and 55 µM, respectively.

Encouraged by these results, we next performed a systematic study of phase separation of Arg/Gly-rich polypeptides and ssRNA rU_40_ as a function of salt concentration and temperature, which represents the most stringent and most comprehensive benchmark currently available for heterotypic condensation^53^. We considered five peptide variants derived from the RGRGG motif, including (RGPGG)_5_, (RPRPP)_5_, (RGRGG)_5_, (RGFGG)_5_, and (RGYGG)_5_ (referred to as RPG, RRP, RRG, RFG, and RYG, respectively), which were experimentally designed to modulate the stability and material properties of the peptide-RNA condensate. As summarized in **Fig. 4d**, direct HyRes/iConNA phase co-existence simulations reproduced the overall phase behavior, including both temperature/salt dependence and relative stability of peptide variants, with only modest deviations in the phase boundaries. This highly nontrivial result underscores the transferability and well-balanced treatment of diverse interactions within the integrated HyRes/iConNA framework. We next asked whether the simulation could capture condensate material properties beyond phase boundaries. At 303 K and 150 mM NaCl, we calculated the monomeric diffusion coefficients (*D*) of both peptides and RNA within condensates (**Extended Data Fig. 3d**). Comparison with experimental condensate viscosities (*η*), measured from the diffusion of 200-nm polystyrene particles^53^, revealed a fractional Stokes-Einstein relationship, *D* ∝ *η*^*α*^, with *α* ≈ 0.2 for both peptide and RNA (**Fig. 4e**). The corresponding correlations between log(D) and log(*η*) are strong with *r* ~ −0.98 and −0.91, respectively, providing an important validation of material properties predicted by HyRes/iConNA simulations.

The simulations also revealed that the peptide-to-RNA ratio within condensates consistently exceeded what is expected from simple charge neutralization (**Fig. 4f** and **Extended Data Fig. 3d**), with the strongest peptide enrichment observed for RFG and RYG. This finding indicates that interactions beyond electrostatic interactions, such as π-π and cation-π interactions, contribute substantially to condensate formation. The RRG-RNA system further illustrates how these interactions shape phase behavior across salt concentrations(**Fig. 4d** and **4g)**. At low NaCl concentrations, both RRG and RNA partitioned into the condensate, with a peptide-to-RNA concentration ratio of approximately 4:1 (**Fig. 4g**, left). Increasing the NaCl concentration above 800 mM progressively screened electrostatic interactions and eliminated phase separation. At 1200 mM NaCl, however, condensation re-emerged and was accompanied by a marked change in the condensate composition. The dense phase is now dominated by RGG, whereas only a small number of RNA molecules accumulate at the droplet surface (**Fig. 4g**, right, and **in Fig. 4d**). Thus, the high-salt condensate represents a distinct peptide-rich phase driven by Arg-Arg interactions, consistent with previous experimental observations^54–56^.

We further tested the HyRes/iConNA framework for heterotypic protein/DNA phase separation by simulating Arg/Lys-rich peptide (pL22) with ssDNA10 and dsDNA10 (see **Fig. 4h** for sequences). The results show that iConDNA reproduces the dynamic random coil conformation of ssDNA10 and the stable duplex structure of dsDNA10 (**Fig. 4h**). Furthermore, the direct phase coexistence simulations accurately reproduced the phase boundaries of both pL22-ssDNA10 and pL22-dsDNA10 mixtures (**Fig. 4i-j** and **Extended Data Fig. 3e-f**), with only modest deviations at elevated NaCl concentrations^57^. Collectively, these stringent benchmarks demonstrate that the integrated HyRes/iConNA framework provides a well-balanced and transferable description of both intra- and intermolecular interactions across proteins and nucleic acids, establishing a versatile platform for quantitative modeling of heterotypic protein-nucleic acid phase separation.

### Extension to cellular metabolites: ML-guided CG mapping and parameterization

Biomolecular condensates can selectively segregate, concentrate, and modulate biochemical activities of metabolites in the cell^17,58^, and metabolites in turn can act as regulators or even drivers of condensate formation and function^59,60^. To extend the HyRes/iConNA framework toward this broader cellular context, we developed a systematic approach for incorporating metabolites and other small molecules. Given the large chemical diversity, we developed a ML protocol, iConMapper, to automatically construct the CG representation for any new chemical matter following the mapping scheme established for HyRes and iConNA (**Fig. 5a**). For this, we retrained the deep supervised graph partitioning model (DSGPM)^61^ with the CG type prediction (DSGPM-TP)^62^ using a pre-annotated dataset of 1000 common metabolites selected from the 3392 detected and quantified ones in the Human Metabolome Database (HMDB)^63^ (see **Methods** and **Extended Data Fig. 4a**). iConMapper assigns the atomistic-to-CG mapping scheme and predicts the best CG bead types (**Extended Data Fig. 4b**). This is followed by all-atom simulations to parameterize bonded terms to accurately capture the covalent geometry and (implicit) dynamics, consistent with the HyRes/iConNA convention (see **Methods**). Using this protocol as implemented in the SMILES-to-ITP, we are constructing a HyRes/iConNA-compatible intermediate resolution library, named iConMetabolome, which includes ~100 members currently and is continuously expanded as additional molecules are incorporated. A representative subset of the resulting metabolite models is shown in **Fig. 5b**.

**Fig. 5.**
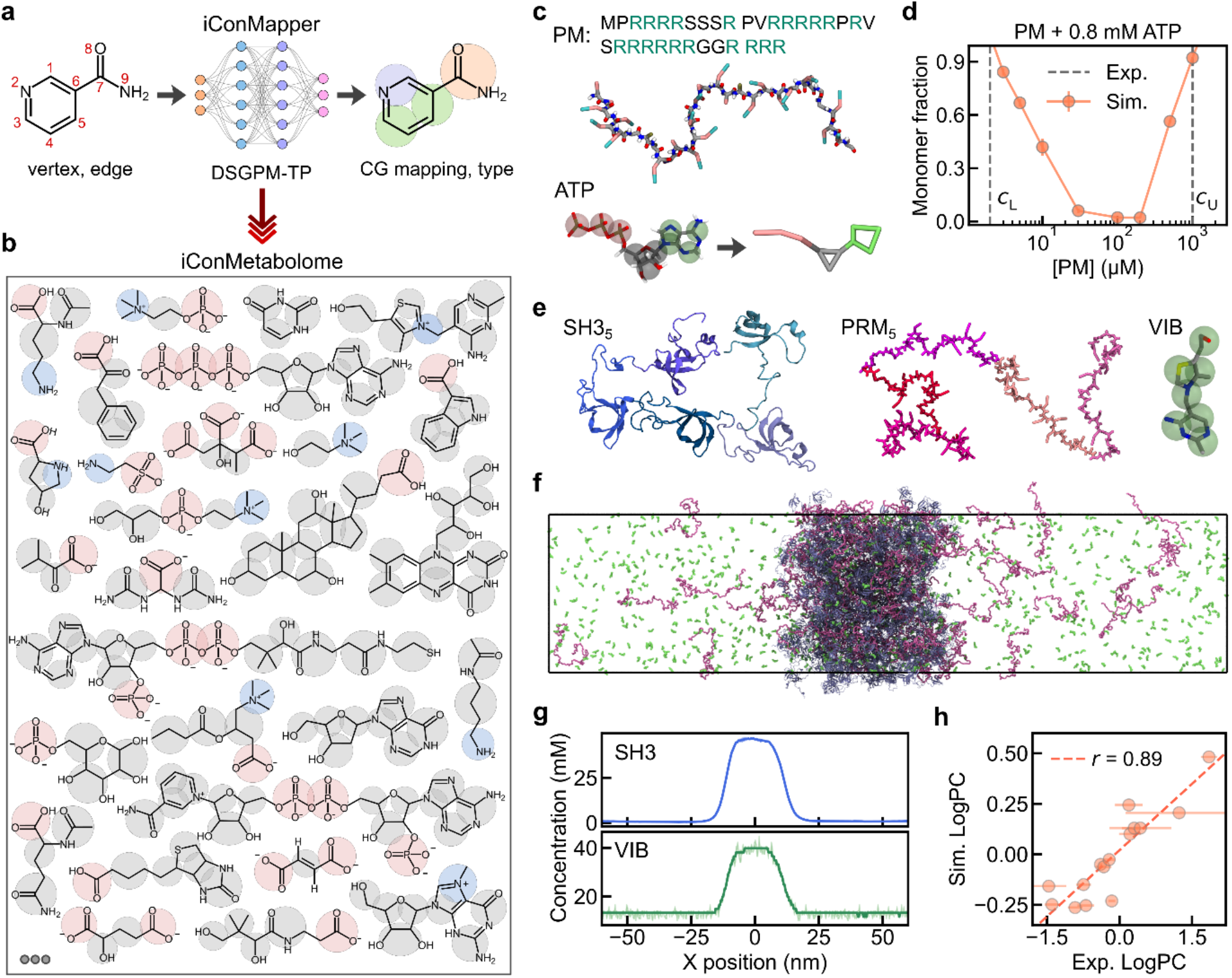
iConMetabolome for simulation of metabolite-condensate interactions. **a**, Schematic workflow of iConMapper. Molecular structures are represented as graphs consisting of vertices and edges, which are processed by the DSGPM-TP model trained to predict the CG mapping and bead types. **b**, Representative subset of the iConMetabolome library. Positively and negatively charged beads are shown in red and blue, respectively. **c**, Sequence and representative snapshot of the PM peptide, together with the CG model of ATP. **d**, Phase separation of PM-ATP mixtures characterized by the monomer fraction. Higher monomer fractions indicate weaker phase separation. The lower and upper boundaries of the experimental phase diagram are indicated by grey dashed lines. **e**, Representative snapshots of SH3_5_, PRM_5_, and vitamin B1 (VIB). The five SH3 and PRM repeats are shown in similar colors. **f**, Representative snapshot of an SH3_5_-PRM_5_ condensate and the corresponding molecular partitioning, with SH3_5_, PRM_5,_ and VIB shown in ice blue, mauve, and lime, respectively. **g**, Density profiles of SH3 and VIB along the *x-*axis, showing enrichment of VIB within condensates. **h**, Comparison of simulation-derived and experimental metabolite partitioning coefficients (LogPC) for 15 metabolites.

Metabolites can critically regulate phase equilibrium of biomolecular condensates. For example, adenosine triphosphate (ATP) is widely involved in modulating protein phase separation^64–66^. We first evaluated iConMetabolome by simulating ATP-induced phase separation of protamine (PM; **Fig. 5c**)^66^. At 0.8 mM ATP, PM undergoes phase separation at sub-µM concentrations, with an experimental critical concentration of ~2 µM^66^. Our simulations showed that the phase behavior depended strongly on the PM-to-ATP ratio, consistent with the experimental observations^66^. Notably, the simulations reproduced both the lower and upper critical PM concentrations delimiting the phase-separated regime. At 3 µM PM, ~80% of PM remained monomeric, although small clusters were still observed (**Fig. 5d** and **Extended Data Fig. 4c**). At 1 mM, over 90% of PM molecules remained monomeric, with only small oligomers detected (**Extended Data Fig. 4c**). We further examined the composition of the condensates across different PM concentrations. The PM-to-ATP ratio within condensates remained nearly constant at ~0.2 across the conditions examined (**Extended Data Fig. 4d**), close to the charge-neutralization ratio of 0.19 for the PM-ATP mixtures, indicating the dominant role of charge-charge interactions in PM and ATP phase transitions.

Biomolecular condensates give rise to distinct local biochemical environments, which can either enrich or exclude metabolites, thereby regulating their biochemical activities^8^. We further evaluated the performance of iConMetabolome in predicting the partitioning of metabolites into condensate^15^. We chose the well-defined condensate formed by the SRC homology 3 (SH3) domain and its proline-rich motif (PRM) ligand^2^ as the host system, and selected 15 metabolites (**Supplementary Table S5**) with partitioning coefficients (PC) spanning LogPC ~ −1.5 to 1.9. We first tested the phase separation of polySH3 and polyPRM (SH3_5_ and PRM_5_; **Fig. 5e**). Under the experimental conditions (10 µM SH3_5_ and PRM_5_), the droplet containing 80 SH3_5_ and 80 PRM_5_ remained stable over the 5-µs simulation, while exhibiting dynamic exchange with the dilute phase (**Extended Data Fig. 4e**). To better sample metabolite partitioning, we constructed a slab configuration (see **Methods**; **Fig. 5f**). PC was directly calculated from the distribution of metabolites across the dense and dilute phases; for example, vitamin B (VIB) molecules were highly enriched within the condensate (**Fig. 5g**). Across the test set of 15 metabolites (**Extended Data Fig. 4f**), HyRes/iCon captured the experimentally measured partitioning trend^15^ with a correlation of *r* ~ 0.89. We note that the range of simulation-derived LogPC values is considerably narrower than the experimental reference. This discrepancy may arise from two factors: smooth CG potentials and reduced degrees of freedom, which weaken entropic and enthalpic specificity, and the absence of context-dependent interactions to faithfully capture the crowded environment of the condensates^7,19^.

## Discussion

Biomolecular condensates operate within the complex cellular environment, where proteins, nucleic acids, ions and/or metabolites engage in dynamic interactions that collectively govern condensate formation, dynamics, composition, properties and thus function. Computational modeling and simulation of these heterogeneous multi-component systems necessarily rely on simplified molecular representations. A central challenge has been to retain sufficient molecular detail, such that one can faithfully capture balance of diverse physical forces while maintaining the computational efficiency required for accessing the time and length scales. Here, we bridge this gap by developing and integrating intermediate-resolution models of proteins, nucleic acids, and metabolites into a unified physics-based platform for quantitative simulation of biomolecular condensates at near-cellular complexity.

The intermediate resolution of the HyRes/iCon framework proves optimal for describing key physical biomolecular interactions important for dynamic proteins and nucleic acids and their phase transitions. The atomistic backbone of HyRes enables explicit treatment of backbone hydrogen bonding and accurate description of transient secondary structures that are pervasive in disordered proteins and segments and important in their phase behaviors. This is illustrated by Dsk2, where HyRes captures the three transient amphipathic helices within the disordered regions and their dynamic engagement with the STI1 domain, which have been shown experimentally to drive Dsk2 condensate formation yet fundamentally inaccessible to single bead models. The six-to seven-bead-per-nucleotide resolution of iConNA allows explicit description of phosphate electrostatics, base stacking, base pairing, and divalent ion-mediated effects, enabling the model to fold RNA nanostars from random coils, capture their kissing-loop-mediated dimerization and condensation, and reproduce the experimentally measured fractal dimension of their condensate networks. Equally important is the self-consistent integration of HyRes and iConNA, where cross-interactions are derived from a principled calibration grounded in all-atom free energy calculations. A single unified parameter set quantitatively reproduces the complex salt- and temperature-dependent phase diagrams of diverse protein–RNA and protein–DNA systems, along with condensate material properties, without the need for system-specific tuning. This shows clearly the transferability and balance of the underlying interaction potentials in the HyRes/iConNA framework.

We have further extended HyRes/iCon to include metabolites through iConMetabolome, such that we can model how condensates can selectively concentrate or exclude metabolites and drug molecules in cells, establishing distinct biochemical microenvironments for function. The ML-enabled SMILES-to-ITP pipeline provides a scalable method for generating consistent CG models of small molecules. Benchmarking showed that the resulting models reproduce experimental condensate partitioning trends across 15 selected metabolites with a correlation of ~0.89. While the predicted dynamic range of partitioning coefficients remains compressed, reflecting the smoothness of coarse-grained potentials, introducing explicit context-dependent interactions will likely improve quantitative accuracy. Several other areas of improvement exist toward modeling true cellular complexity. Firstly, additional post-translational and post-transcriptional modifications are needed, which are key regulators of condensates. Secondly, strategies are needed to model non-equilibrium processes such as condensate aging and liquid-to-solid transitions relevant to neurodegenerative disease. Last but not least, as new experimental studies emerge, systematic benchmarking against increasingly heterogeneous mixtures is needed, rather than relying on the mostly binary and ternary systems.

Importantly, the ability of the current HyRes/iCon framework to reproduce a wide range of experimental observables strongly support its viability as a platform for generating mechanistic hypotheses that can guide new experiments. Quantitative simulations of biomolecular condensates towards cellular complexity provide crucial molecular-level descriptions of transient structures, dynamic interaction networks, and compositional heterogeneity within condensates, allowing one to address questions that remain inaccessible to experiments. For example, how specific interaction motifs contribute to condensate specificity, how RNA structure modulates phase behavior and material properties, and how metabolite partitioning influences enzymatic activity within condensates. As the field increasingly moves from identifying and characterizing phase-separating systems toward understanding how condensate properties are tuned for function in cells and become dysregulated in disease, we anticipate that the quantitative HyRes/iCon simulation platform will play a central role.

## Methods

### Integration of HyRes/iCon: balancing protein-nucleic acid interactions

Details of the HyRes and iConNA energy functions are given in the **Supplementary Text**. Integration of HyRes and iCon requires three main modifications. First, the dielectric constant in iConRNA was increased from the original value of 20 to 60 (at room temperature) to be consistent with HyRes. Second, HyRes was revised to follow the empirical temperature-dependent dielectric constant used in iConNA,^67^

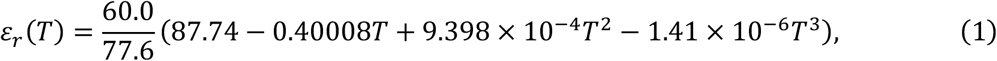

where *T* is the temperature in °C. The pre-factor 60.0/77.6 was introduced to scale the dielectric constant such that *ε*_r_ = 60.0 at 303 K (30 °C). Both HyRes and iConRNA were then systematically recalibrated independently, as done previously^37,39^.

The third and most important modification is the balancing of protein-nucleic interactions given the recalibrated HyRes and iConNA models. For this, we introduced specific pairwise interaction potentials to better capture π-π, cation-π, amide-π, and strong hydrogen bond interactions between selected amino acid side chains, including Arg, Lys, Asp, Glu, Asn, Gln, Phe, Tyr, and Trp, and nucleobases, where standard Lennard-Jones potentials using the combination rule are insufficient. We first assigned the relative strengths among these interactions based on pairwise potentials of mean force derived from all-atom simulations^50^. All interaction strengths were then scaled based on that of the Arg-Arg interaction, which has already been calibrated in the HyRes protein model^37^. The strengths of all specific pairwise interactions are summarized in **Supplementary Table S6**. For computational efficiency and consistency with the CG representation, a single representative bead was selected from each aromatic ring to mediate these interactions (see **Supplementary Table S6**). For Arg and Lys, the specific pairwise interactions were imposed on the charged beads (**Fig. 1a**).

### HyRes/iCon simulation protocols

The HyresBuilder package (https://github.com/lslumass/HyresBuilder) was developed for preparation and simulation of all systems in conjunction with OpenMM^68^. All single-chain simulations were performed under non-periodic boundary conditions. Initial conformations were generated using HyresBuilder, followed by energy minimization and equilibration. Production simulations were performed for 4 µs, with the final 2 µs used for analysis. For MDPs, folded domains were treated as rigid bodies to maintain the folded structures obtained from crystal structures or the AlphaFold database^69^. The folded domains included in the simulations are listed in **Supplementary Table S2**. The initial configurations of phase transition simulations were prepared using Packmol^70^, given the relaxed monomer structures generated by single-chain equilibration simulations. Cubic periodic boxes were used in all phase co-existence simulations (see below). All simulations were performed using Langevin dynamics with a collision frequency of 0.1 ps^-1^ and an integration time step of 8 fs. All bonds involving hydrogen atoms were constrained using the SHAKE algorithm^71^. Nonbonded interactions were smoothly cut off from 1.0 to 1.2 nm for Lennard-Jones interactions and from 1.6 to 1.8 nm for Debye-Hückel electrostatic interactions.

To estimate the melting temperatures of RNA hairpins and duplexes, stable folded structures were first obtained from folding simulations at 273 K. The RNA molecules were then simulated at temperatures ranging from 30 to 85°C in 5°C increments, following the single-chain simulation protocol above. For the 14-bp RNA duplex, a higher temperature range of 50-95 °C was used. The number of base pairs formed at each temperature was used to calculate the folded fraction, defined as the ratio of the instantaneous number of base pairs to the theoretical number. The melting temperature was determined as the temperature at which the folded fraction reached 0.5 (**Supplementary Fig. S2**).

### Data analysis

Trajectory analysis was performed using MDAnalysis^72^ and an in-house Python package (SciKit, https://github.com/lslumass/SciKit). Unless otherwise stated, convergence was assessed using block analysis (two blocks), and error bars represent the standard deviations across blocks. Cluster analysis of phase co-existence simulations was performed based on residue-residue distances between Cα atoms for proteins and phosphate (P) groups for nucleic acids. Two chains were assigned to the same cluster when at least one interchain residue pair was separated by less than 8 Å. This procedure was used to track the number of monomers, number of clusters, and cluster sizes throughout the trajectories. Protein condensate contact maps were calculated using Cα and all side chain beads, with a contact cutoff of 6 Å. A total of 1000 frames were uniformly extracted from the final 1 µs of each trajectory for contact analysis. Secondary structure assignments and scattering profiles were calculated using DSSP^73^ and Pepsi-SAXS^74^, respectively. VMD^75^ was used for visualization and rendering snapshots. Charge pattern (*κ*)^76^ was estimated using CIDER^77^.

### Phase-coexistence simulations for saturation concentration determination

Direct-coexistence simulations were performed to determine *C*_sat_ in HyRes. Initial droplets were prepared by packing protein molecules into a large cubic box, followed by compaction using a short 10-ns NPT simulation at 1 bar. The resulting droplets were then transferred to NVT simulation boxes of appropriate sizes, which were initially determined based on the experimental *C*_sat_ such that approximately 10% of the molecules would remain in the dilute phase. The box size was adjusted for systems when few monomers remained in the dilute phase or the droplet largely dissolved. The simulations were performed for 4-10 µs, as required for convergence. The simulation box sizes, the final configurations, and simulation lengths are summarized in the **Supplementary Table S7**, with representative snapshots shown in **Supplementary Fig. S3a**. The number of monomers in the dilute phase was determined from the equilibrated trajectories (**Supplementary Fig. S3b-c**), with the uncertainty estimated as the standard deviation between the two equal blocks. The droplet radius was obtained by fitting the radial density profile to (**Supplementary Fig. S3b**)

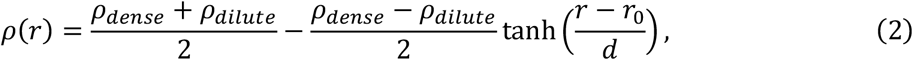

where *ρ*_dense_ and *ρ*_dilute_ are the densities of the dense and dilute phases, respectively. *r*_0_ is the droplet radius, and *d* is the interfacial width. *C*_sat_ was then calculated as *N*_dilute_/(*V*_box_ − *V*_droplet_), where *N*_dilute_ is the number of proteins in the dilute phase, and *V*_box_ and *V*_droplet_ are the volumes of the simulation box and the droplet, respectively.

### Construction and simulation of the Dsk2 condensate

The initial structure of Dsk2 was predicted using AlphaFold^69^, followed by a 10-µs single-chain simulation using the protocol above for MDPs to fully equilibrate the disordered segments and organization of the folded domains. A snapshot was selected from the equilibrated trajectory (**Fig. 2d**) and used as the conformation for constructing the initial simulation box using Packmol^70^. The simulation box contains 100 copies of Dsk2; the cubic box was eventually relaxed to a size of 50 nm in each dimension. All folded domains were treated as rigid bodies, consistent with the single-chain MDP simulation protocol. The final NVT production simulation lasted 4 µs following 2 µs equilibration simulation. Simulation convergence was evaluated by comparing the density profiles and intermolecular contact maps between two 2-µs blocks.

### RNA nanostars folding, dimerization, and phase separation

The initial ssRNA single strand was built as a random coil using HyresBuilder and sampled at 298 K following the single-chain simulation protocol. Folding occurred within the first few hundred nanoseconds, and the nanostar structures remained stable for the rest of the 2 µs simulation (**Fig. 3d**). A folded snapshot was randomly selected to serve as the initial nanostar conformation for preparing the phase separation simulations. 100 copies of folded 3sWT or 3sv5 nanostars were randomly distributed in a 120-nm cubic box (100 µM), followed by minimization and equilibration, and then subjected to 4-µs production runs at 298 and 333 K. Cluster analysis showed that the largest cluster (a 40-mer) remained stable over the final 1 µs of the simulation. A short 500-ns simulation of dilute RNA nanostars (100 copies of 3sWT in a 255-nm cubic box, equivalent to 10 µM) was also performed to examine the ability of the kissing loop to drive dimerization.

### Determination of heterotypic protein/nucleic acid phase diagrams

To construct the phase diagram of the peptide/nucleic acid phase separation, we performed a series of simulations to probe the estimated phase boundary, instead of attempting to achieve phase co-existence equilibrium directly. The latter requires excessive simulations due to the much slower protein/nucleic acid exchange between the dilute and condensed phases. For example, to determine the NaCl concentration below which 100 µM rU_30_/(PR)_30_ can undergo phase separation (**Fig. 4a**), we first simulated the experimentally measured transition point (300 mM NaCl) for 2 to 4 µs to detect droplet formation (**Supplementary Fig. S4a**). If large clusters formed under this condition, we tested progressively higher NaCl concentrations (400 mM, 500 mM, and beyond) until droplet formation was no longer observed; if no large cluster formed, we tested progressively lower NaCl concentrations. This approach allowed the phase boundary to be estimated with a small number of modest simulations, which are summarized in **Supplementary Tables S8-10**.

After at least 2 µs of simulation, the final phase boundary was determined from several complementary metrics (**Supplementary Fig. S4**). We first measured the size of the largest cluster to quickly identify a candidate transition point (**Supplementary Fig. S4b**), which we then confirmed using the number of remaining monomers in the simulation box (**Supplementary Fig. S4c**). The time evolution of monomer numbers was fit to a biexponential function,

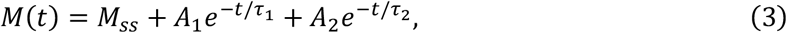

where *M*(*t*) and *M*_ss_ are the number of monomers at time *t* and at the steady-state plateau, respectively, and *τ* = (*A*_1_*τ*_1_ + *A*_2_*τ*_2_)/(*A*_1_ + *A*_2_) is the relaxation time. Far from the transition, the dispersed state is thermodynamically favored, and the system relaxes quickly to steady state, yielding a short relaxation time. Conversely, deep within the phase-separated regime, cluster formation is strongly favored, and under these supersaturated conditions monomers are rapidly and decisively incorporated into clusters, again giving a short relaxation time. Only near the transition, where the system has no net thermodynamic preference between monomeric and clustered states, does the relaxation time become large. The peak in the *τ* plotted against [NaCl] (or other conditions) therefore indicates the transition point (**Supplementary Fig. S4d**). This estimate was cross-checked against structural analysis using the static structure factor

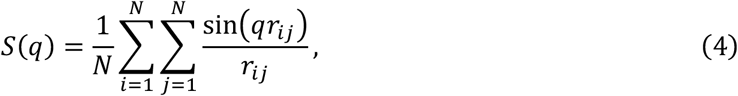

where *N* is the number of peptide and/or nucleic acid chains, whose centers of mass (COM) were used to compute the structure factor. *r*_*ij*_ is the pairwise distance between COM *i* and *j*, and *q* is the wavenumber. *S*(*q*) was fit to the Ornstein-Zernike form (**Supplementary Fig. S4e**)

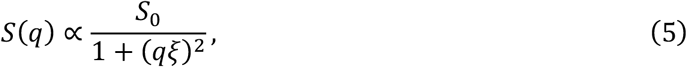

where *S*_0_ and *ξ* are the extrapolated zero-wavevector structure factor and correlation length, respectively, describing the amplitude and length scale of density fluctuations. Near the transition, growing long-wavelength density fluctuations cause *ξ* to reach a maximum while *S*_0_ rises sharply (**Supplementary Fig. S4f** and **g**). For the same example of 100 µM rU_30_/(PR)_30_, these analyses together indicated that the critical NaCl concentration for phase transition lies between 400 and 500 mM. We used 400 mM as an approximate phase boundary in the final phase diagram (**Fig. 4a**).

### Training of iConMapper for automatic all-atom-to-iCon mapping

iConMapper was developed by retraining the DSGPM-TP model^62^. We first manually labeled 1000 metabolites following the CG mapping and typing convention of HyRes and iConNA to build the training dataset (https://github.com/lslumass/iConMetabolome), spanning a broad range of chemical classes. This dataset size has been reported as sufficient for effective training in prior work (~1100 and 400 molecules, respectively)^61,62^. In the DSGPM framework, molecules are converted to graphs encoding vertex and edge information, including bond degrees, cycle/aromatic structures, and charges. In our training, the charge feature was muted. We retrained the model on this dataset using a 5-fold cross-validation scheme, with hidden and output embedding dimensions of 128. Adaptive moment estimation (Adam) was used for optimization (learning rate = 1e-3, no weight decay, batch size = 32) for 800 epochs per fold, minimizing a weighted sum of a triplet-cut loss (margin = 1.0, weight = 1.0) and a CG-type loss (weight = 0.5)^62^. The triplet-cut loss optimizes the partitioning of atoms into CG beads, and the CG-type loss optimizes the CG bead type label. For each fold, the best-performing checkpoint on the test set was retained, and final results were averaged across the best epochs from all five folds.

### Parameterization of bonded terms in iCon metabolite models

Besides iConMapper automatic assignment of iCon CG types, simulation of a new metabolite requires parameters for bonded terms of the iCon CG metabolite models. Unless previously reported for metabolites with identical CG mapping^78^, we performed all-atom simulations to obtain reference distributions of pseudo bonds, angles, and dihedrals. These simulations were prepared using CHARMM-GUI^79^ with default simulation settings and performed using GROMACS 2025^80^ with the OpenFF force field^81^ at 293.15 K for 500 ns. Because bonded parameters can differ substantially between molecules even when the corresponding two CG bead types are identical, we assigned and stored bonded parameters separately for each metabolite. This molecule-specific parameter table is implemented in HyresBuilder to facilitate automated construction of CG metabolite models. The resulting reference distributions for 32 metabolites examined in this work are summarized in **Supplementary Fig. S5-9**.

The iCon framework currently includes full parameterization for ~100 metabolites (e.g., **Supplementary Table S11**), with additional metabolites incorporated on an ongoing basis. All associated bonded parameters have been implemented in HyresBuilder. A Python wrapper, Smiles2itp.py, has been developed for integration of custom small molecules. Given the SMILES string, the script generates the itp parameter file (see **Supplementary Text** for an example) following a standard protocol: 1) convert the SMILES into a CG model (CG structure and mapping scheme) using iConMapper; 2) perform a 500-ns all-atom simulation of the small molecule in vacuum or water using GROMACS with either OpenFF or GAFF2 (and AM1-BCC partial charges) force field; and 3) derive the CG bonded parameters by fitting reference distributions from the all-atom trajectory and generate the corresponding itp file. The resulting parameter file can be automatically incorporated into HyresBuilder and ready for simulation within the HyRes/iConNA framework. Although the SMILES-to-ITP pipeline is fully automated, manual inspection of the CG mapping scheme and the agreement between the all-atom and fitted CG distributions are recommended before performing simulations.

### Simulation of PM-ATP phase separation

PM and ATP molecules were first equilibrated for 1 µs at 298 K, from which random snapshots were selected as their initial structures for the following phase separation simulations. PM and ATP molecules were mixed to a reference concentration based on the experimental ratio. Details can be found in **Supplementary Table S12**. 10 µs production simulations were performed for each condition, and the final 2 µs were used for analysis.

### Metabolite partitioning in the SH3_5_/PRM_5_ condensate

We first confirmed phase separation of SH3_5_ and PRM_5_. Following the direct-coexistence simulation protocol, a droplet containing 80 SH3_5_ and 80 PRM_5_ chains was prepared and relaxed in a 240-nm cubic box (10 µM each). After brief minimization and equilibration, a 5-µs production simulation was performed, during which the droplet remained stable (**Extended Data Fig. 4e**). To sample the partitioning of small molecules into the condensate, we used a slab configuration (**Fig. 5f**). Here, 80 SH3_5_ and 80 PRM_5_ chains were first pre-compacted in a cubic box with a side length of 28 nm, and the box was then expanded to 120 nm along the *x*-axis. The system was subsequently relaxed and equilibrated for 5 µs. To quantify metabolite partitioning, 1000 copies of each metabolite were randomly distributed throughout the slab simulation box. After at least 500 ns of equilibration, the spatial distributions of each metabolite along the *x*-axis were calculated and fit to the corresponding density profile,

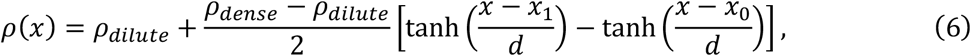

where *x*_1_ and *x*_2_ denote the midpoints of the left and right interfaces, respectively. The partition coefficient (LogPC) was calculated as

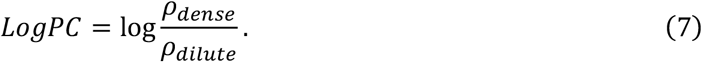

## Acknowledgements

This work is supported by NIH through R35 GM144045 and by NSF through CHE 2516941. The computational resources for this work were provided by the University of Massachusetts Amherst’s partnership with the Unity Research Computing Platform, a multi-institutional cluster led by the University of Massachusetts and the University of Rhode Island.

## Author contributions

Li and Chen conceptualized the idea. Li conducted HyRes/iConRNA refinement and benchmarks. Ng and Li designed and performed simulations and analysis of iConDNA. Li and Chen drafted and revised the manuscript with inputs from Ng.

## Additional information

Extended data is available for this paper at https://doi.org/xxxx

## Supplementary information

The online version contains supplementary material available at https://doi.org/xxxx

---

**Extended Data Fig 1.**
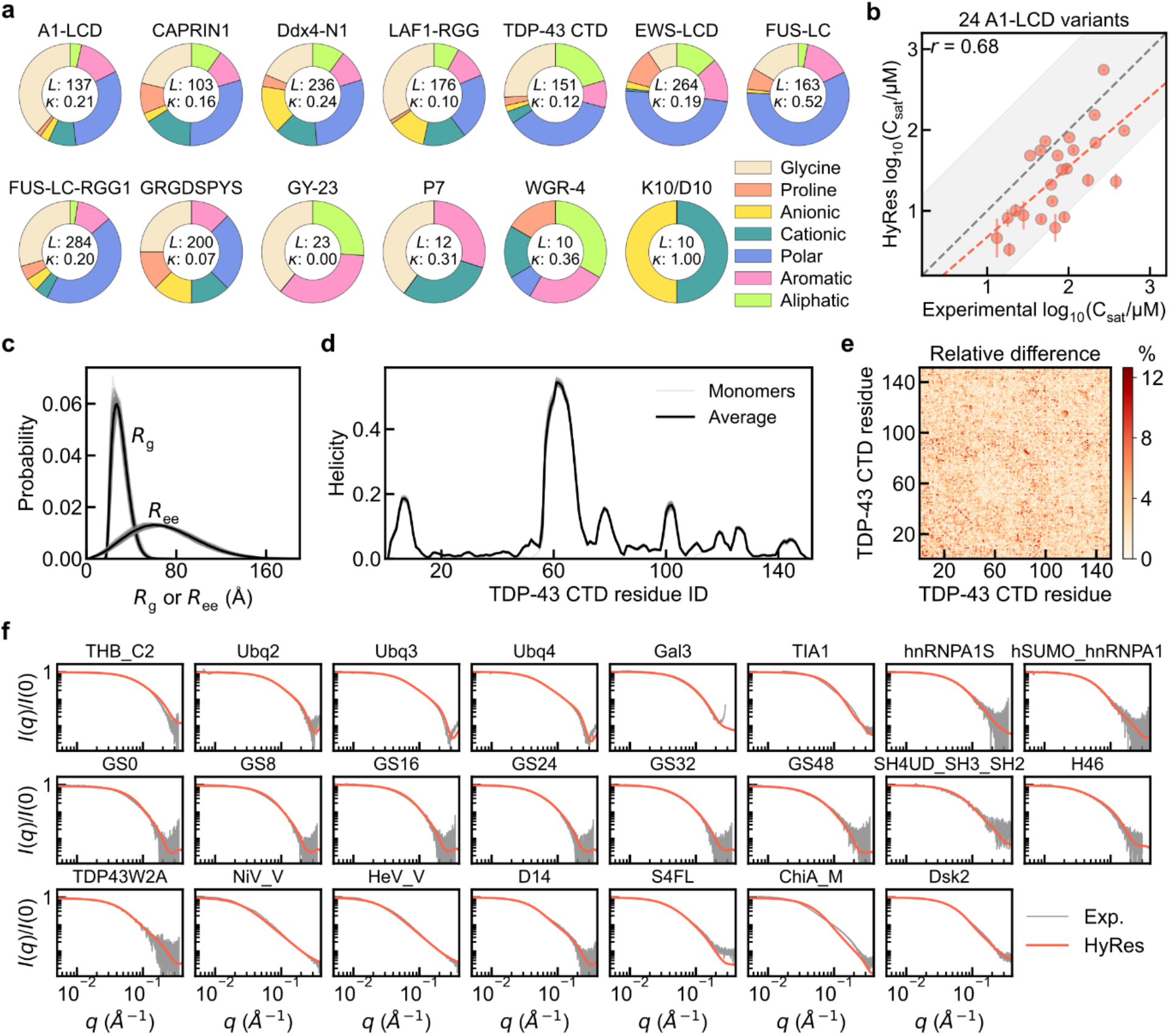
HyRes accurately and efficiently captures protein phase equilibrium. **a**, Pie charts showing the diverse sequence compositions of the 13 WT IDPs. The sequence length (*L*) and charge pattern (*κ*) are indicated. **b**, Comparison of HyRes-derived and experimental *C*_sat_ for 24 A1-LCD variants. The gray shaded region indicates values within 10-fold of the experimental values. Grey dashed diagonal lines plot *y* = *x*, and red dashed lines plot the best linear fit. Error bars in the calculated values were estimated from block analysis. **c**, *R*_g_ and *R*_ee_ distributions and **d**, residue helicity profiles of each of the 100 TDP-43 monomers within the simulation box. In **c** and **d**, monomeric distributions are shown in grey lines, with their averages shown in black lines. **e**, Percent deviations between the two intermolecular residue-residue contact probability maps within the condensate of TDP-43 LCD derived from the first and second halves of the last 2 µs of the trajectory. **f**, Comparison between HyRes-derived and experimental SAXS profiles of 23 MDPs (see **Supplementary Table S2**).

**Extended Data Fig 2.**
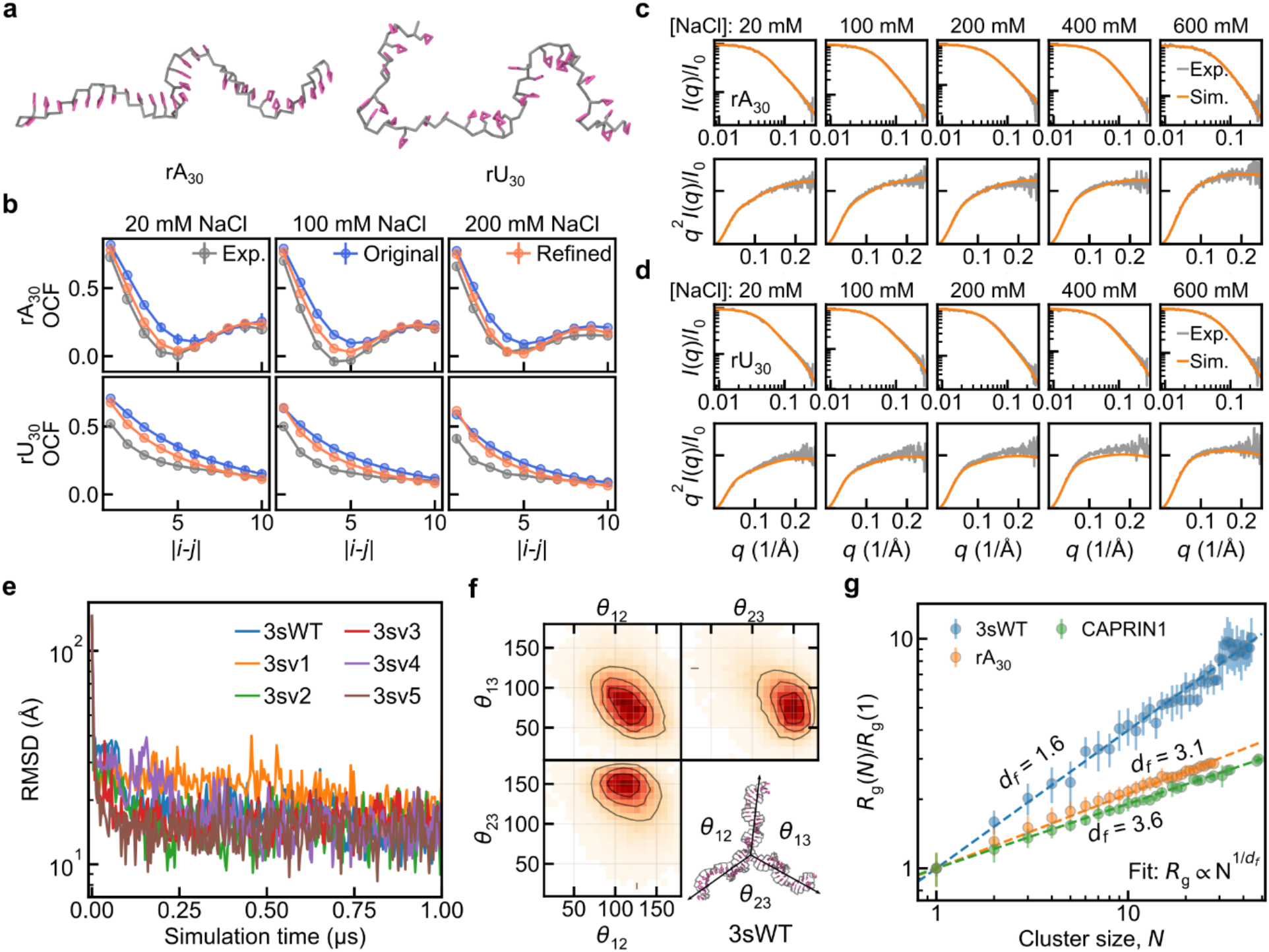
Revised iConRNA and phase separation of RNA nanostars. **a**, Representative snapshots of rA_30_ and rU_30_. Backbones and bases are colored in grey and mauve, respectively. **b**, Comparison of orientation correlation functions (OCFs) of rA_30_ and rU_30_ under various NaCl concentrations from experiments, original and refined iConRNA models. **c,d**, Comparison between HyRes-derived and experimental SAXS profiles of rA_30_ (**c**) and rU_30_ (**d**), respectively, under various NaCl concentrations. Lower rows show Krathy plots. **e**, Folding of 6 nanostar variants indicated by the decrease in RMSD to the average structures. **f**, Conformational ensembles of 3swt nanostar shown through the combination of the angles between every two arms. **g**, Fractal scaling of cluster *R*_g_ as a function of the number of RNA molecules (*N*) within the cluster for the condensates of 3swt, rA_30_, and CAPRIN1. The dashed line shows the fit of *R*_g_ ∝ *N*^1/*d*f^, where *d*_f_ is the fractal dimension. Error bars indicate the range of *R*_g_ for the same *N*-size cluster.

**Extended Data Fig 3.**
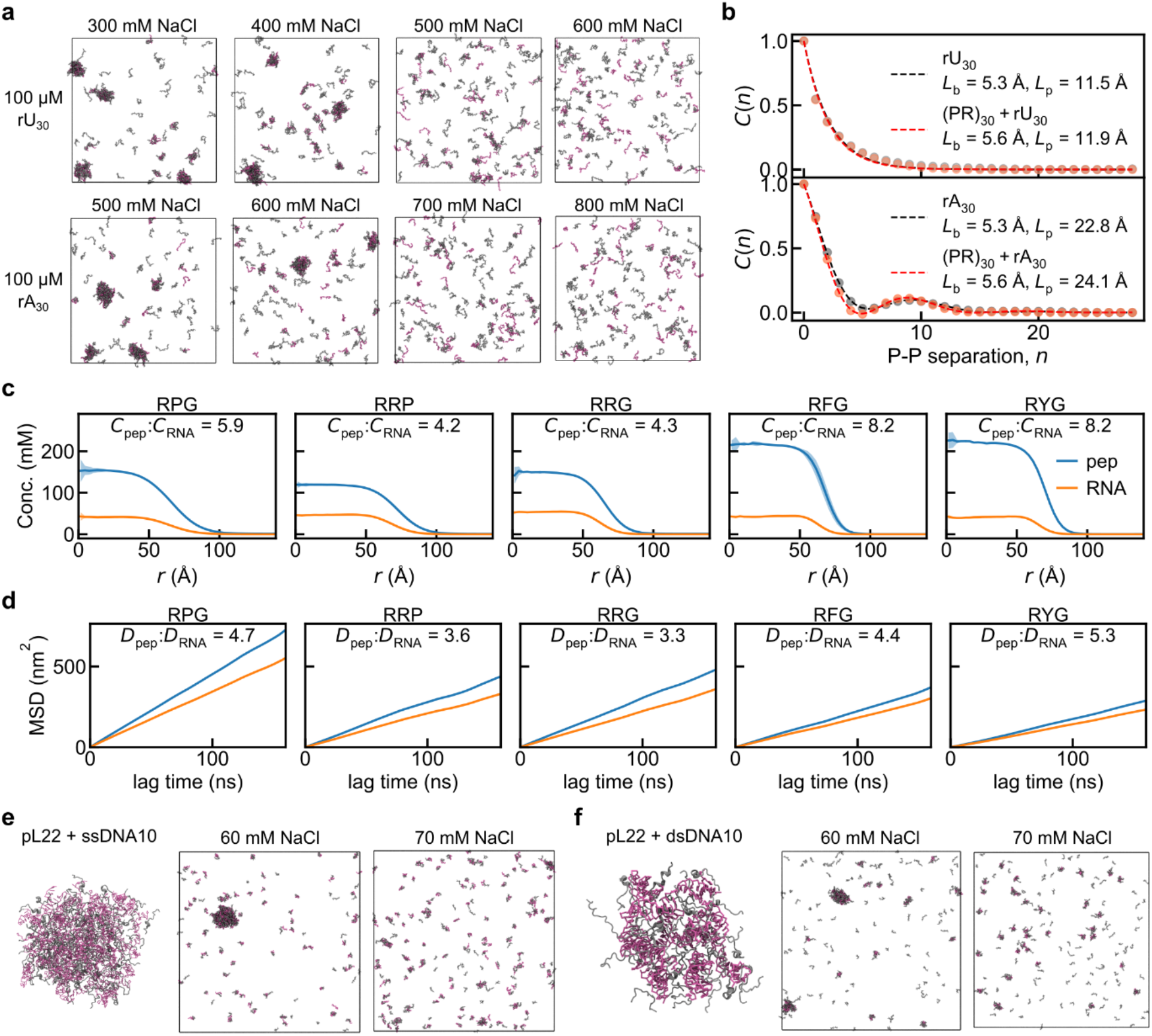
HyRes/iConNA simulations of the heterotypic phase separation of proteins and nucleic acids. **a**, Representative snapshots for the phase separation of (PR)_30_-rU_30_ and (PR)_30_-rA_30_ systems under different salt conditions. **b**, Calculation of persistence lengths by fitting the backbone orientation correlation function using the HWLC model (see **Methods**). **c**, Radial density profiles of various peptide-RNA condensates, with the concentration ratio between peptide and RNA within condensates noted. **d**, MSD of peptide and RNA within various peptide-RNA condensates, with the ratio of protein and RNA diffusion constants shown. **e,f**, Representative snapshots for the droplets and phase coexistence of pL22+ssDNA10 (**e**) and pL22+dsDNA (**f**) systems under two salt conditions. Peptide and DNA molecules are shown in grey and mauve, respectively.

**Extended Data Fig 4.**
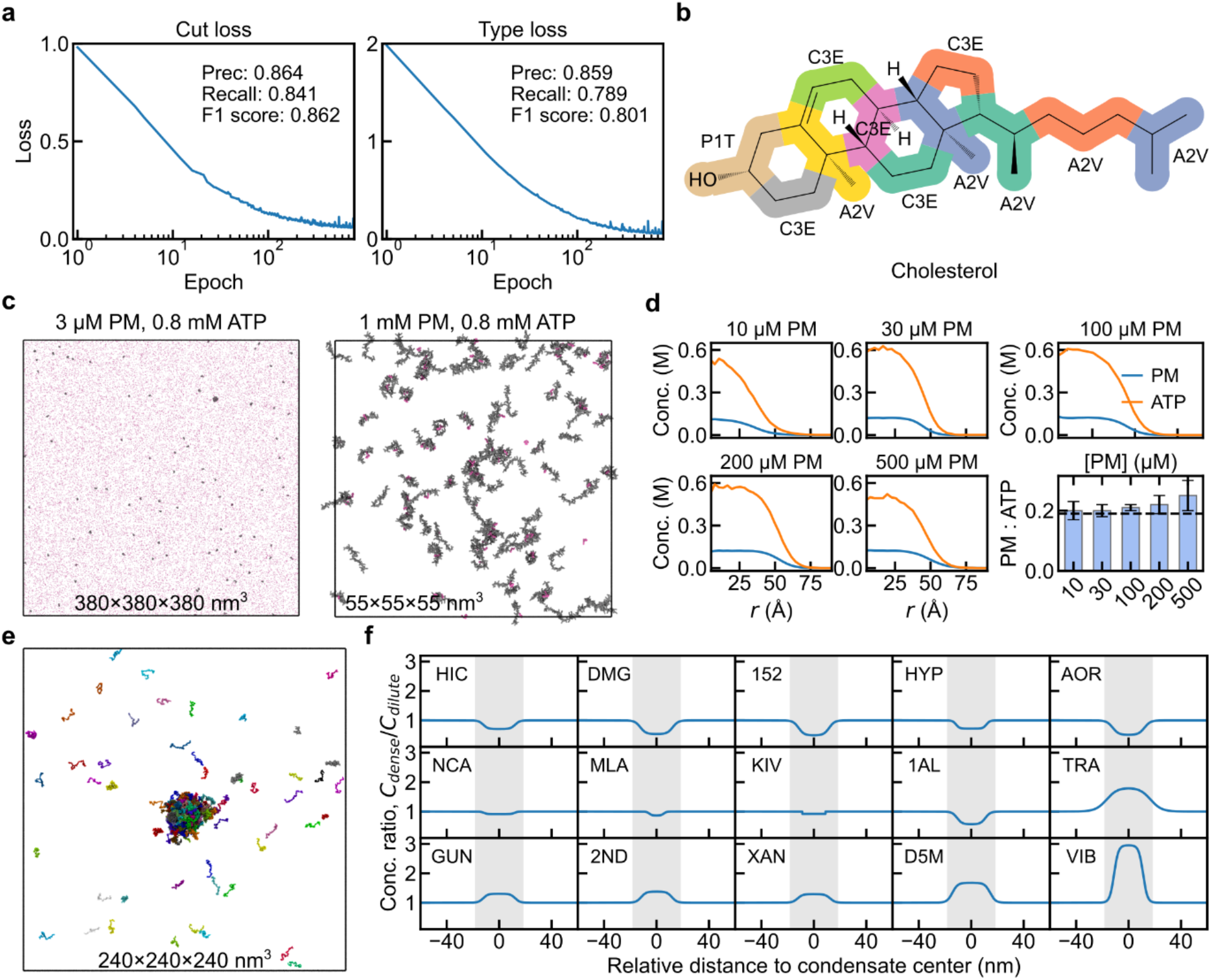
Training and performance of iConMetabolome. **a**, Validation cut loss and type loss, as well as the precision, recall, and F1 score for iConMapper. The cut loss and type loss are contributed by CG bead mapping and corresponding bead type. **b**, Example of iConMapper assignment of CG beads and types for cholesterol. Colored groups indicate the CG bead mapping with predicted bead types assigned. **c**, Final snapshots for the simulation of mixtures of 3 µM and 1 mM PM together with 0.8 mM ATP. **d**, Radial density profiles (in M) of PM and ATP for various PM concentrations. The ratios of PM to ATP are summarized in the last panel. Error bars are estimated from block analysis. **e**, Representative snapshot for the SH3_5_-PMR_5_ condensate, derived from the direct-coexistence simulation of 80 SH3_5_ and 80 PMR_5_ (10 µM each). **f**, Distributions of metabolites in the slab simulations. Concentrations were normalized as the ratio of dense to dilute concentrations (*C*_dense_/*C*_dilute_). Condensates were highlighted as shadowed regions.

## Supplementary Information

### Supplementary Text

#### HyRes/iCon energy function

The HyRes/iCon framework includes HyRes protein, iConRNA, and iConDNA models, and the additional iConMetabolome library for small molecules. The total potential includes

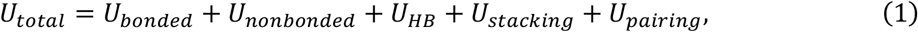

where the right terms are bonded, nonbonded, hydrogen bond, stacking, and pairing interactions. *U*_bonded_ contains contributions from the standard bond, angle, dihedral angle, and improper dihedral angle terms, which are

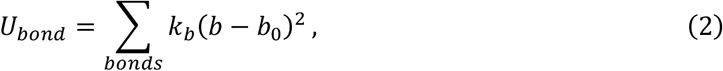

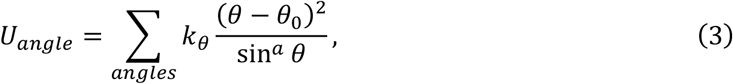

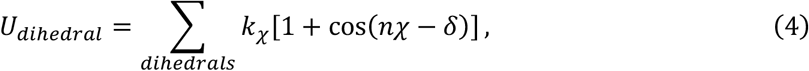

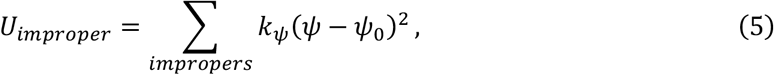

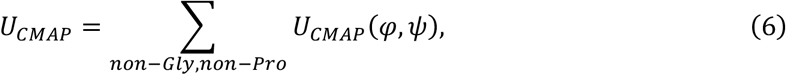

For angle terms, *a* = 1 is used for angles in protein backbones, meaning a normal harmonic angle potential, while *a* = 2 for other angles, meaning a restricted bending (ReB) potential^1^.

Nonbonded interactions *U*_*nonbonded*_ include both electrostatic and Van der Waals terms,

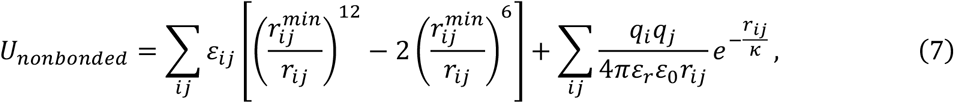

where 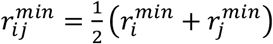, and 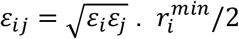 and *ε*_*i*_ are the vdW radius and interaction strength of bead *i. q*_*i*_ is the charge (−1 for B1 and 0 otherwise), and *r*_*ij*_ is the distance between beads *i* and *j*. *ε*_0_ is the permittivity of vacuum. *κ* is the Debye screening length, determined as 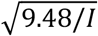 (in Å) at 300 K, where *I* is the ionic strength in molarity. *ε*_*r*_ is the effective dielectric constant, which is set to 60 at 300 K.

*U*_*HB*_ is specific to backbone hydrogen bonds of proteins to describe transient structures of IDPs. The potential form is

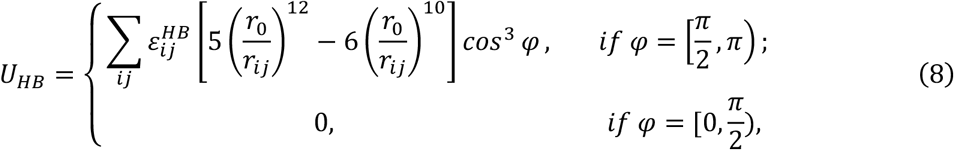

where 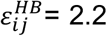 kcal/mol is the strength of the hydrogen bond, *r*_*ij*_ is the N-O distance with a cutoff distance of 0.45 nm and an equilibrium distance (*r*_0_) of 0.29 nm. *φ* is the angle of N-H-O with a cutoff of 90 degrees.

For nucleic acids, explicit base stacking was described as

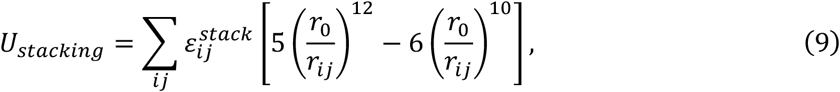

where |*i* − *j*| = 1, and 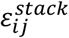 is the stacking strength between bases *i* and *j*. The explicit base pairing was described as

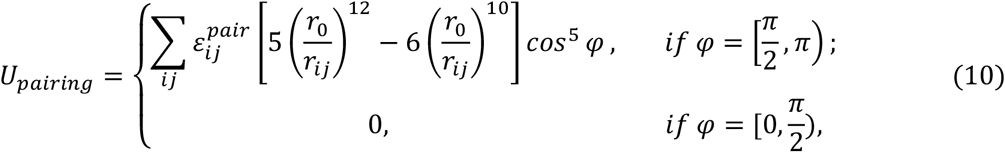

where 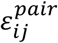 is the pairing strength, *r*_*ij*_ is the distance between corresponding CG beads. *φ* is the angle formed by two paired bases, which is used to roughly mimic the planar structure of two paired bases. *r*_0_ is the equilibrium distance for each pair of hydrogen-bonding CG beads. More details can be found in the original iConRNA^2^ and iConDNA models.

#### Example of an itp file for custom small molecules

To define a custom small molecule compatible with the iCon framework, an itp file should be provided following the format below.

~~~
[ RESI ] CKC
[ ATOM ]
M1 A1F 0.00
M2 A2F 0.00
M3 A3F 0.00
[ BOND ]
M1 M2 334.0 2.22
M1 M3 340.0 2.18
M2 M3 328.0 2.22
[ ANGL ]
M1 M2 M3 51.0 103.0
[ DIHE ]
[ IMPR ]
~~~

### Supplementary Tables

**Supplementary Table S1.** IDPs for the benchmark of *C*_sat_ (µM) prediction.

| IDPs | Exp. $C_{\text{sat}}$ | Sim. $C_{\text{sat}}$ | IDPs | Exp. $C_{\text{sat}}$ | Sim. $C_{\text{sat}}$ |
| --- | --- | --- | --- | --- | --- |
| A1-NLS3 | 102.20 | 80.63 | FUS-LC_12S-A4 | 171.82 | 50.18 |
| A1+NLS3 | 73.40 | 48.29 | FUS-LC_12S-N4 | 258.40 | 44.69 |
| A1-10R3 | 382.00 | 23.09 | FUS-LC_12S-Q4 | 78.46 | 71.23 |
| A1-6R3 | 45.20 | 55.72 | FUS- RGG14 | 13.51 | 61.84 |
| A1+2R3 | 95.50 | 33.19 | FUS- RGG1_S-G4 | 4.32 | 2.96 |
| A1+7R3 | 173.60 | 23.92 | FUS- RGG1_T-S4 | 10.89 | 36.50 |
| A1-3R+3K3 | 83.10 | 32.20 | FUS- RGG1_Q-A4 | 34.43 | 1.16 |
| A1-6R+6K3 | 483.70 | 98.21 | FUS- RGG1_Q-G4 | 13.63 | 7.50 |
| A1-2K3 | 13.20 | 4.61 | FUS- RGG1_Q-N4 | 39.9 | 8.33 |
| A1-12F+12Y3 | 61.10 | 21.13 | FUS- RGG1_Q-S4 | 30.91 | 52.92 |
| A1+7F-7Y3 | 209.10 | 153.68 | GY235 | 364.0 | 981.31 |
| A1-8F+4Y3 | 63.20 | 13.16 | GY23-F17A6 | 1000.0 | 1760.79 |
| A1-9F+3Y3 | 115.00 | 56.31 | GY23-F9A6 | 720.0 | 2107.70 |
| A1+12E3 | 214.80 | 69.54 | GY23-F9L7 | 494.0 | 1914.89 |
| A1-4D3 | 88.80 | 8.36 | GY23-A7F7 | 110.0 | 456.55 |
| A1+4D3 | 17.90 | 8.17 | GY23-2AF7 | 12.40 | 150.12 |
| A1+8D3 | 18.70 | 3.31 | GY23-3AF7 | 6.20 | 61.10 |
| A1+7R+12D3 | 33.70 | 48.18 | GY25-V15 | 500.0 | 330.81 |
| A1+7K+12D3 | 270.40 | 554.47 | GY25-V25 | 700.0 | 3802.04 |
| A1+23G-23S3 | 46.10 | 7.89 | GY205 | 350.0 | 1107.32 |
| A1-10G+10S3 | 28.00 | 8.81 | GRGDSPYS8 | 3.1 | 6.61 |
| A1-20G+20S3 | 52.00 | 72.67 | EWS-LCD9 | 25.0 | 22.31 |
| A1-14N+14Q3 | 22.40 | 10.08 | Ddx4-N110 | 23.8 | 102.57 |
| A1-23S+23T3 | 68.70 | 6.19 | LAF1-RGG11 | 44.2 | 65.32 |
| FUS-LC4 | 126.46 | 183.07 | LAF1-RGGs11 | 5.5 | 3.51 |
| FUS-LC_S-G4 | 33.35 | 3.74 | LAF1-RGGsp11 | 3.4 | 6.67 |
| FUS-LC_T-S4 | 115.39 | 123.55 | LAF1-RGGd11 | 321.5 | 160.57 |
| FUS-LC_4QQ-GG4 | 127.17 | 110.90 | TDP-43 CTD12 | 13.7 | 60.31 |
| FUS-LC_4QQ-AA4 | 180.18 | 110.15 | K10-D1013 | 3.7 | 36.19 |
| FUS-LC_4QQ-NN4 | 178.29 | 120.98 | WGR-413 | 7480.0 | 3040.0 |
| FUS-LC_4QQ-SS4 | 178.14 | 303.70 | P714 | 3280.0 | 12840.0 |
| FUS-LC_12S-G4 | 125.45 | 79.81 | CAPRIN115 | 980.0 | 1020.0 |

**Supplementary Table S2.**
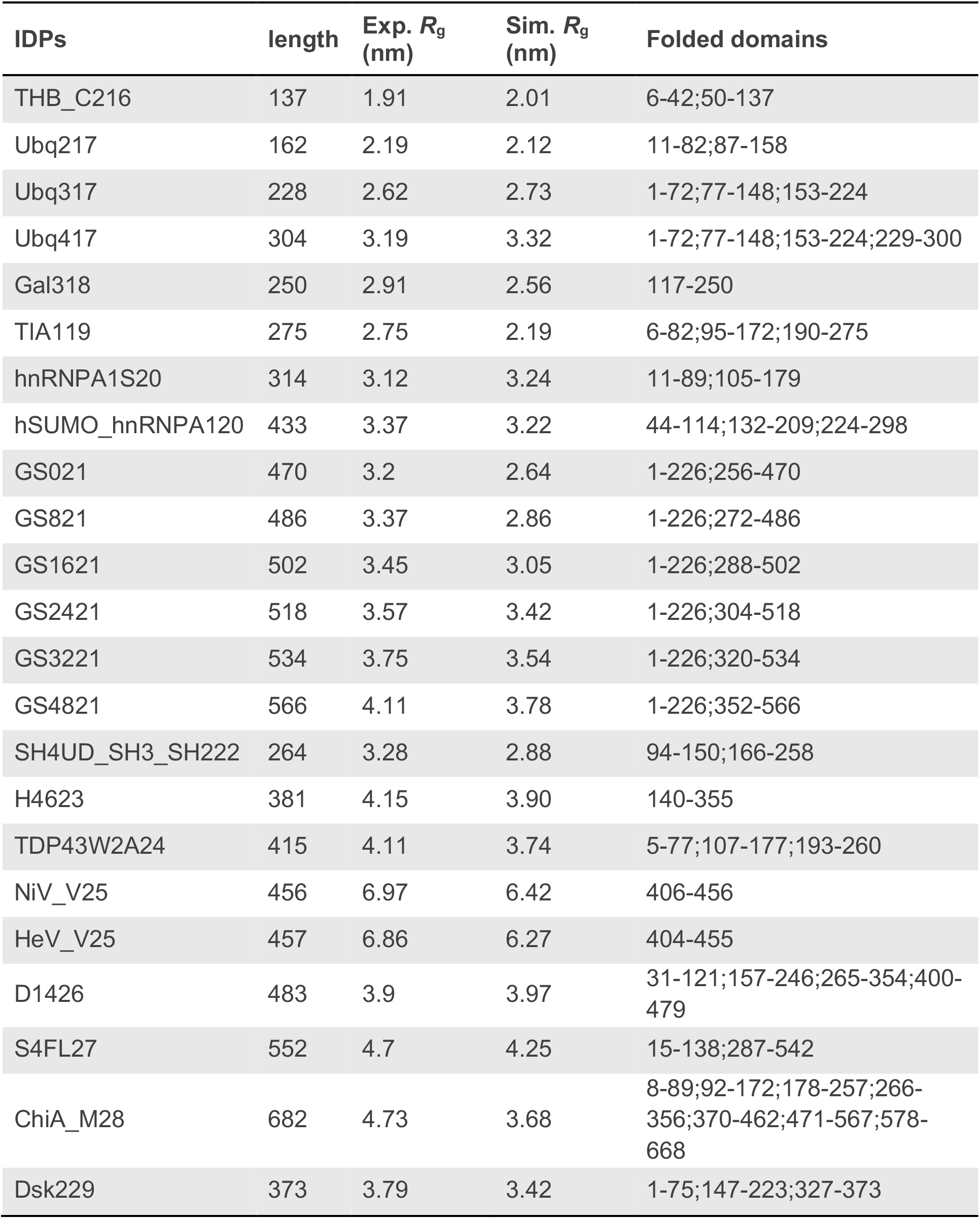
MDPs used in the benchmark.

| IDPs | length | Exp. $R_g$<br>(nm) | Sim. $R_g$<br>(nm) | Folded domains |
| --- | --- | --- | --- | --- |
| THB_C216 | 137 | 1.91 | 2.01 | 6-42;50-137 |
| Ubq217 | 162 | 2.19 | 2.12 | 11-82;87-158 |
| Ubq317 | 228 | 2.62 | 2.73 | 1-72;77-148;153-224 |
| Ubq417 | 304 | 3.19 | 3.32 | 1-72;77-148;153-224;229-300 |
| Gal318 | 250 | 2.91 | 2.56 | 117-250 |
| TIA119 | 275 | 2.75 | 2.19 | 6-82;95-172;190-275 |
| hnRNPA1S20 | 314 | 3.12 | 3.24 | 11-89;105-179 |
| hSUMO_hnRNPA120 | 433 | 3.37 | 3.22 | 44-114;132-209;224-298 |
| GS021 | 470 | 3.2 | 2.64 | 1-226;256-470 |
| GS821 | 486 | 3.37 | 2.86 | 1-226;272-486 |
| GS1621 | 502 | 3.45 | 3.05 | 1-226;288-502 |
| GS2421 | 518 | 3.57 | 3.42 | 1-226;304-518 |
| GS3221 | 534 | 3.75 | 3.54 | 1-226;320-534 |
| GS4821 | 566 | 4.11 | 3.78 | 1-226;352-566 |
| SH4UD_SH3_SH222 | 264 | 3.28 | 2.88 | 94-150;166-258 |
| H4623 | 381 | 4.15 | 3.90 | 140-355 |
| TDP43W2A24 | 415 | 4.11 | 3.74 | 5-77;107-177;193-260 |
| NiV_V25 | 456 | 6.97 | 6.42 | 406-456 |
| HeV_V25 | 457 | 6.86 | 6.27 | 404-455 |
| D1426 | 483 | 3.9 | 3.97 | 31-121;157-246;265-354;400-479 |
| S4FL27 | 552 | 4.7 | 4.25 | 15-138;287-542 |
| ChiA_M28 | 682 | 4.73 | 3.68 | 8-89;92-172;178-257;266-356;370-462;471-567;578-668 |
| Dsk229 | 373 | 3.79 | 3.42 | 1-75;147-223;327-373 |

**Supplementary Table S3.**
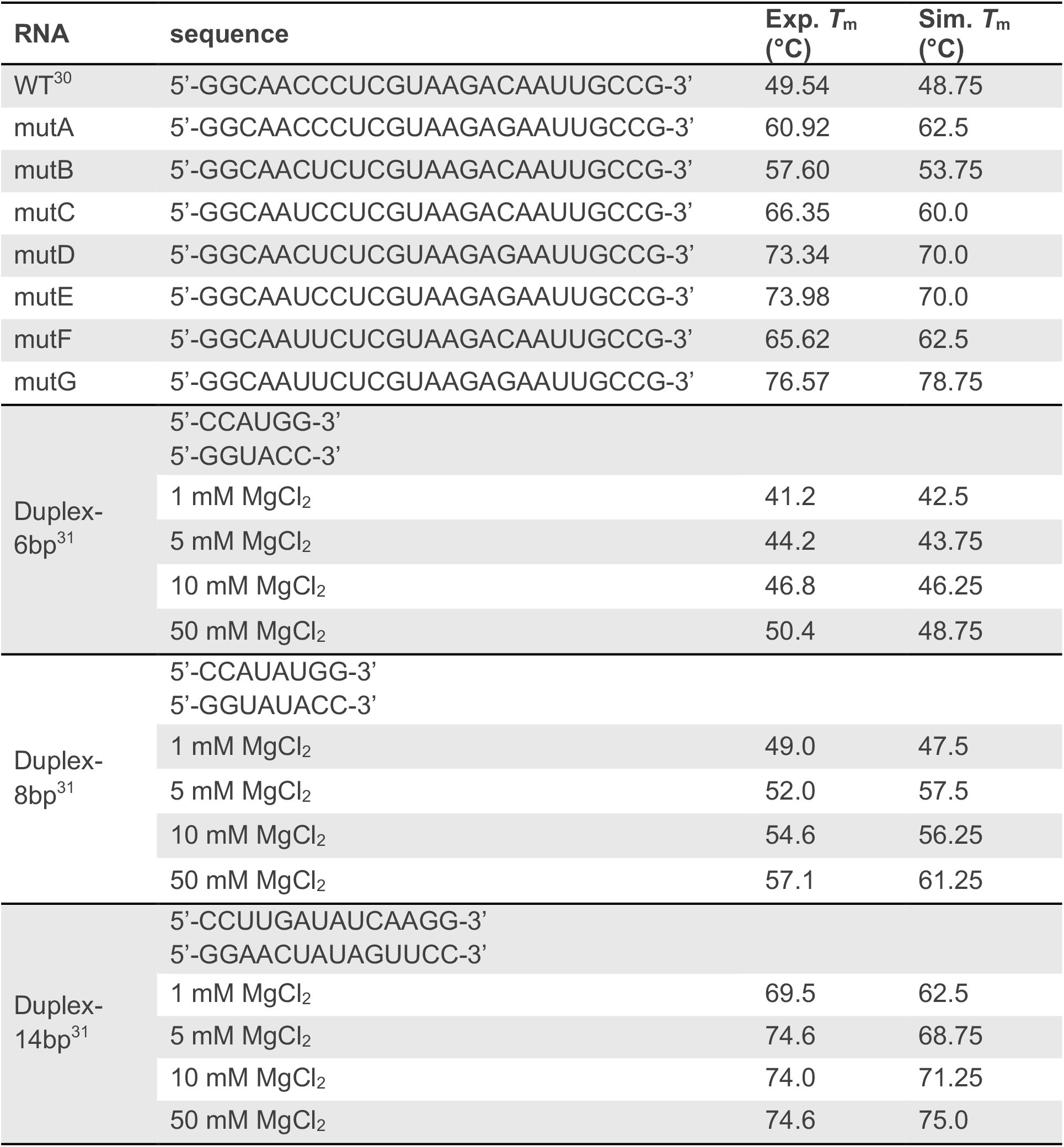
RNA sequences used for the iConRNA benchmark.

**Supplementary Table S4.**
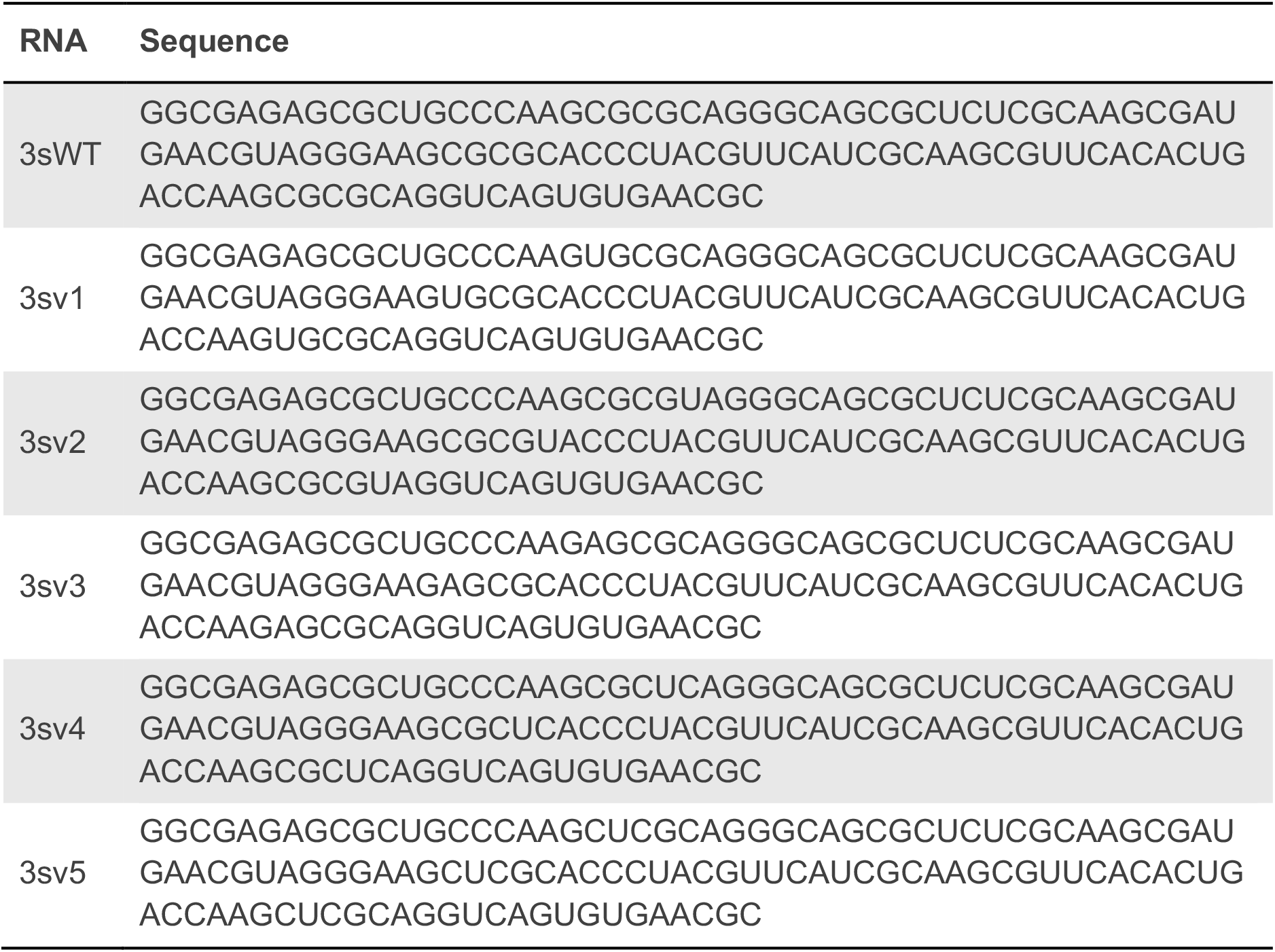
Sequences of RNA nanostars^32^.

**Supplementary Table S5.** 15 metabolites and their partitioning coefficients^33^.

| Metabolite | 3-character CCD | Exp. LogPC | Sim. LogPC |
| --- | --- | --- | --- |
| 1-Methyl-Histidine | HIC | -1.464 | -0.158 |
| Dimethylglycine | DMG | -1.397 | -0.249 |
| Carnitine | 152 | -0.919 | -0.264 |
| Hydroxyproline | HYP | -0.744 | -0.150 |
| N-acetyl-L-ornithine | AOR | -0.696 | -0.254 |
| Nicotinamide | NCA | -0.393 | -0.050 |
| Malonic acid | MLA | -0.326 | -0.062 |
| 2-keto-isovalerate | KIV | -0.214 | -0.026 |
| Allantoate | 1AL | -0.160 | -0.231 |
| cis-Aconitate | TRA | 0.193 | 0.244 |
| Guanine | GUN | 0.217 | 0.101 |
| Deoxyinosine | 2ND | 0.304 | 0.130 |
| Xanthine | XAN | 0.432 | 0.130 |
| dAMP | D5M | 1.234 | 0.206 |
| Thiamine | VIB | 1.851 | 0.482 |

**Supplementary Table S6.** Amino acid-nucleotide interaction matrix^34^. Pairwise interactions were assigned to the CG bead pairs to mimic the cross-interactions.

| Interaction pair<br>(CG beads) | | PMF<br>(kcal/mol) | $\epsilon$<br>(kcal/mol) | Interaction pair | | PMF<br>(kcal/mol) | $\epsilon$<br>(kcal/mol) |
| --- | --- | --- | --- | --- | --- | --- | --- |
| ARG<br>(QdR) | TYR<br>(P1Y) | -4.42 | -0.65 | ARG<br>(QdR) | CYT<br>(RC1) | -2.53 | -0.38 |
| ARG<br>(QdR) | PHE<br>(A1F) | -3.35 | -0.50 | ARG<br>(QdR) | URA<br>(RU1) | -0.63 | -0.09 |
| LYS<br>(QdK) | TYR<br>(P1Y) | -2.04 | -0.30 | ASP<br>(QaD) | ADE<br>(RA1) | -0.63 | -0.09 |
| LYS<br>(QdK) | PHE<br>(A1F) | -2.02 | -0.30 | ASP<br>(QaD) | GUA<br>(RG1) | -1.73 | -0.26 |
| TYR<br>(A2Y) | ADE<br>(RA1) | -6.07 | -0.86 | ASP<br>(QaD) | CYT<br>(RC1) | -2.06 | -0.31 |
| TYR<br>(A2Y) | GUA<br>(RG1) | -6.75 | -0.96 | ASP<br>(QaD) | URA<br>(RU1) | -1.41 | -0.21 |
| TYR<br>(A2Y) | CYT<br>(RC1) | -4.63 | -0.65 | GLU<br>(QaE) | ADE<br>(RA1) | -1.24 | -0.19 |
| TYR<br>(A2Y) | URA<br>(RU1) | -6.08 | -0.86 | GLU<br>(QaE) | GUA<br>(RG1) | -2.87 | -0.43 |
| PHE<br>(A2F) | ADE<br>(RA1) | -5.57 | -0.78 | GLU<br>(QaE) | CYT<br>(RC1) | -0.36 | -0.06 |
| PHE<br>(A2F) | GUA<br>(RG1) | -5.58 | -0.78 | GLU<br>(QaE) | URA<br>(RU1) | -0.66 | -0.10 |
| PHE<br>(A2F) | CYT<br>(RC1) | -5.18 | -0.72 | GLN<br>(P4Q) | ADE<br>(RA1) | -0.77 | -0.11 |
| PHE<br>(A2F) | URA<br>(RU1) | -5.40 | -0.76 | GLN<br>(P4Q) | GUA<br>(RG1) | -3.00 | -0.45 |
| LYS<br>(QdK) | ADE<br>(RA1) | -0.17 | -0.05 | GLN<br>(P4Q) | CYT<br>(RC1) | -3.01 | -0.45 |
| LYS<br>(QdK) | GUA<br>(RG1) | -1.50 | -0.22 | GLN<br>(P4Q) | URA<br>(RU1) | -2.51 | -0.41 |
| LYS<br>(QdK) | CYT<br>(RC1) | -1.81 | -0.27 | ASN<br>(P5N) | ADE<br>(RA1) | -2.74 | -0.41 |
| LYS<br>(QdK) | URA<br>(RU1) | -1.16 | -0.17 | ASN<br>(P5N) | GUA<br>(RG1) | -2.94 | -0.44 |
| ARG<br>(QdR) | ADE<br>(RA1) | -0.87 | -0.13 | ASN<br>(P5N) | CYT<br>(RC1) | -2.30 | -0.34 |
| ARG<br>(QdR) | GUA<br>(RG1) | -2.85 | -0.43 | ASN<br>(P5N) | URA<br>(RU1) | -1.67 | -0.25 |

**Supplementary Table S7.** Simulation details of the phase co-existence simulations, including number of monomers (*N*_m_), box length (*L*_box_), temperature (*T*), NaCl concentration (*C*_s_), and simulation time (*t*), for calculating *C*_sat_ of various IDPs in HyRes.

| IDPs | $N_m$ | $L_{\text{box}}$<br>(nm) | $T$<br>(K) | $C_s$<br>(mM) | $t$<br>(μs) | IDPs | $N_m$ | $L_{\text{box}}$<br>(nm) | $T$<br>(K) | $C_s$<br>(mM) | $t$<br>(μs) |
| --- | --- | --- | --- | --- | --- | --- | --- | --- | --- | --- | --- |
| A1-NLS | 100 | 50 | 293 | 150 | 10 | 12S-A | 100 | 50 | 297 | 150 | 10 |
| +NLS | 100 | 60 | 293 | 150 | 10 | 12S-N | 100 | 40 | 297 | 150 | 10 |
| -10R | 100 | 40 | 289 | 150 | 10 | 12S-Q | 100 | 60 | 297 | 150 | 10 |
| -6R | 100 | 60 | 293 | 150 | 10 | FUS-RGG1 | 100 | 100 | 297 | 150 | 5 |
| +2R | 100 | 50 | 289 | 150 | 10 | S-G | 100 | 100 | 297 | 150 | 5 |
| +7R | 100 | 50 | 277 | 150 | 10 | T-S | 100 | 100 | 297 | 150 | 5 |
| -3R+3K | 100 | 60 | 277 | 150 | 10 | Q-A | 100 | 70 | 297 | 150 | 5 |
| -6R+6K | 100 | 40 | 277 | 150 | 10 | Q-G | 100 | 100 | 297 | 150 | 5 |
| -2K | 100 | 100 | 293 | 150 | 10 | Q-N | 100 | 70 | 297 | 150 | 5 |
| -12F+12Y | 100 | 60 | 293 | 150 | 10 | Q-S | 100 | 70 | 297 | 150 | 5 |
| +7F-7Y | 100 | 40 | 293 | 150 | 10 | GY23 | 200 | 30 | 298 | 100 | 4 |
| -8F+4Y | 100 | 60 | 277 | 150 | 10 | F17A | 200 | 20 | 298 | 100 | 4 |
| -9F+3Y | 100 | 50 | 277 | 150 | 10 | F9A | 200 | 25 | 298 | 100 | 4 |
| +12E | 100 | 40 | 277 | 150 | 10 | F9L | 200 | 30 | 298 | 100 | 4 |
| -4D | 100 | 60 | 277 | 150 | 10 | A7F | 200 | 50 | 298 | 100 | 4 |
| +4D | 100 | 100 | 289 | 150 | 10 | 2AF | 200 | 80 | 298 | 100 | 4 |
| +8D | 100 | 100 | 277 | 150 | 10 | 3AF | 200 | 100 | 298 | 100 | 4 |
| +7R+12D | 100 | 70 | 303 | 150 | 10 | GY25-V1 | 200 | 30 | 298 | 100 | 4 |
| +7K+12D | 100 | 40 | 293 | 150 | 10 | GY25-V2 | 200 | 25 | 298 | 100 | 4 |
| +23G-23S | 100 | 60 | 293 | 150 | 10 | GY20 | 200 | 30 | 298 | 100 | 4 |
| -10G+10S | 100 | 80 | 277 | 150 | 10 | GRGDSPYS | 100 | 90 | 310 | 150 | 10 |
| -20G+20S | 100 | 60 | 277 | 150 | 10 | EWS-LCD | 100 | 80 | 298 | 150 | 10 |
| -14N+14Q | 100 | 80 | 277 | 150 | 10 | Ddx4-N1 | 100 | 80 | 298 | 150 | 10 |
| -23S+23T | 100 | 60 | 277 | 150 | 10 | LAF1-RGG | 100 | 60 | 296 | 150 | 10 |
| FUS-LC | 100 | 50 | 297 | 150 | 10 | LAF1-RGGs | 100 | 100 | 296 | 150 | 10 |
| S-G | 100 | 80 | 297 | 150 | 10 | LAF1-RGGsp | 100 | 100 | 296 | 150 | 10 |
| T-S | 100 | 50 | 297 | 150 | 10 | LAF1-RGGd | 100 | 40 | 296 | 150 | 10 |
| 4QQ-GG | 100 | 50 | 297 | 150 | 10 | TDP-43 CTD | 100 | 100 | 298 | 150 | 10 |
| 4QQ-AA | 100 | 50 | 297 | 150 | 10 | K10-D10 | 400 | 80 | 298 | 25 | 10 |
| 4QQ-NN | 100 | 50 | 297 | 150 | 10 | WGR-4 | 500 | 30 | 298 | 800 | 10 |
| 4QQ-SS | 100 | 50 | 297 | 150 | 10 | P7 | 500 | 30 | 296 | 250 | 10 |
| 12S-G | 100 | 50 | 297 | 150 | 10 | CAPRIN1 | 100 | 30 | 298 | 400 | 10 |

**Supplementary Table S8.** Simulation details, including number of monomers (*N*_m_), box length (*L*_box_), temperature (*T*), NaCl concentration (*C*_s_), and simulation time (*t*), for determining phase diagrams of (PR)_30_-rA_30_ and (PR)_30_-rU_30_ systems.

| systems | $N_m$ | $L_{\text{box}}$<br>(nm) | $T$<br>(K) | $C_s$<br>(mM) | $t$<br>(μs) | IDPs | $N_m$ | $L_{\text{box}}$<br>(nm) | $T$<br>(K) | $C_s$<br>(mM) | $t$<br>(μs) |
| --- | --- | --- | --- | --- | --- | --- | --- | --- | --- | --- | --- |
| (PR)<sub>30</sub><br>rA<sub>30</sub> | 100<br>50 | 203 | 303 | 200 | 4 | (PR)<sub>30</sub><br>rU<sub>30</sub> | 100<br>50 | 203 | 303 | 20 | 4 |
|  |  |  |  | 300 |  |  |  |  |  | 200 |  |
|  |  |  |  | 400 |  |  |  |  |  | 300 |  |
|  |  |  |  | 500 |  |  |  |  |  | 400 |  |
|  |  |  |  | 600 |  |  |  |  |  | 500 |  |
| (PR)<sub>30</sub><br>rA<sub>30</sub> | 100<br>50 | 181 | 303 | 400 | 4 | (PR)<sub>30</sub><br>rU<sub>30</sub> | 100<br>50 | 181 | 303 | 200 | 4 |
|  |  |  |  | 500 |  |  |  |  |  | 300 |  |
|  |  |  |  | 600 |  |  |  |  |  | 400 |  |
|  |  |  |  | 700 |  |  |  |  |  | 500 |  |
|  |  |  |  | 800 |  |  |  |  |  | 600 |  |
| (PR)<sub>30</sub><br>rA<sub>30</sub> | 100<br>50 | 118 | 303 | 500 | 2 | (PR)<sub>30</sub><br>rU<sub>30</sub> | 100<br>50 | 118 | 303 | 200 | 2 |
|  |  |  |  | 600 |  |  |  |  |  | 300 |  |
|  |  |  |  | 700 |  |  |  |  |  | 400 |  |
|  |  |  |  | 800 |  |  |  |  |  | 500 |  |
|  |  |  |  | 900 |  |  |  |  |  | 600 |  |
| (PR)<sub>30</sub><br>rA<sub>30</sub> | 100<br>50 | 94 | 303 | 500 | 2 | (PR)<sub>30</sub><br>rU<sub>30</sub> | 100<br>50 | 94 | 303 | 200 | 2 |
|  |  |  |  | 600 |  |  |  |  |  | 300 |  |
|  |  |  |  | 700 |  |  |  |  |  | 400 |  |
|  |  |  |  | 800 |  |  |  |  |  | 500 |  |
|  |  |  |  | 900 |  |  |  |  |  | 600 |  |
| (PR)<sub>30</sub><br>rA<sub>30</sub> | 100<br>50 | 70 | 303 | 600 | 2 | (PR)<sub>30</sub><br>rU<sub>30</sub> | 100<br>50 | 70 | 303 | 300 | 2 |
|  |  |  |  | 700 |  |  |  |  |  | 400 |  |
|  |  |  |  | 800 |  |  |  |  |  | 500 |  |
|  |  |  |  | 900 |  |  |  |  |  | 600 |  |
|  |  |  |  | 1000 |  |  |  |  |  | 700 |  |

**Supplementary Table S9.** Simulation details, including number of monomers (*N*_m_), box length (*L*_box_), temperature (*T*), NaCl concentration (*C*_s_), and simulation time (*t*), for determining phase diagrams of rU_40_ and model peptides, including RPG, RRP, RRG, RFG, and RYG.

| Peptides | $N_m$<br>(peptide/RNA) | $L_{\text{box}}$<br>(nm) | $T$<br>(K) | $C_s$<br>(mM) | $t$<br>($\mu\text{s}$) |
| --- | --- | --- | --- | --- | --- |
| RPG | 460/43 | 70 | 283 | 100, 200, 300, 400 | 2 |
|  |  |  | 303 | 100, 150, 200, 300 |  |
|  |  |  | 323 | 10, 50, 100, 200 |  |
|  |  |  | 343 | 10, 50, 100, 200 |  |
|  |  |  | 363 | 10, 50, 100 |  |
| RRP | 329/43 | 70 | 283 | 300, 400, 500, 600 | 2 |
|  |  |  | 303 | 150, 300, 350, 400 |  |
|  |  |  | 323 | 200, 250, 300, 400 |  |
|  |  |  | 343 | 100, 200, 250, 300 |  |
|  |  |  | 363 | 100, 200, 300 |  |
| RRG | 407/43 | 70 | 283 | 600, 700, 800, 900, 1000, 1200 | 2 |
|  |  |  | 293 | 1200, 1500 |  |
|  |  |  | 303 | 150, 400, 500, 600, 1200 |  |
|  |  |  | 323 | 300, 400, 500, 600 |  |
|  |  |  | 343 | 200, 300, 400, 500 |  |
|  |  |  | 363 | 100, 200, 300, 400 |  |
| RFG | 414/43 | 70 | 283 | 800, 1000, 1200, 1500 | 2 |
|  |  |  | 293 | 1200, 1500 |  |
|  |  |  | 303 | 150, 700, 800, 900, 1000, 1200 |  |
|  |  |  | 323 | 300, 400, 500, 600 |  |
|  |  |  | 343 | 200, 300, 400, 500 |  |
|  |  |  | 363 | 100, 200, 300, 400 |  |
| RYG | 401/43 | 70 | 303 | 600, 800, 900, 1200, 1500, 1600 | 2 |
|  |  |  | 313 | 1200, 1500 |  |
|  |  |  | 323 | 500, 600, 700, 800 |  |
|  |  |  | 343 | 300, 400, 500, 600, 1200 |  |
|  |  |  | 363 | 100, 200, 300, 400, 500 |  |

**Supplementary Table S10.** Simulation details, including number of monomers (*N*_m_), box length (*L*_box_), temperature (*T*), NaCl concentration (*C*_s_), and simulation time (*t*), for determining phase diagrams of pL22-ssDNA10 and pL22-dsDNA10.

| DNA | $N_m$<br>(peptide/DNA) | $L_{\text{box}}$<br>(nm) | T<br>(K) | $C_s$<br>(mM) | t<br>(μs) |
| --- | --- | --- | --- | --- | --- |
| ssDNA10 | 200/145 | 103 | 283 | 70, 80, 90 | 10 |
|  |  |  | 298 | 40, 50, 60, 70, 80 |  |
|  |  |  | 308 | 50, 60, 70 |  |
|  |  |  | 328 | 40, 50, 60 |  |
|  |  |  | 348 | 30, 40, 50 |  |
|  |  |  | 368 | 20, 30, 40 |  |
| dsDNA10 | 200/35 | 103 | 283 | 60, 70, 80 | 10 |
|  |  |  | 293 | 60, 70, 80 |  |
|  |  |  | 303 | 40, 50, 60 |  |
|  |  |  | 313 | 40, 50, 60 |  |
|  |  |  | 323 | 30, 40, 50 |  |
|  |  |  | 333 | 20, 30, 40 |  |

**Supplementary Table S11.** Selected parameterized metabolites in iConMetabolites.

| Metabolite | Res | Class | Metabolite | Res | Class |
| --- | --- | --- | --- | --- | --- |
| DHAP | 13P | Others | Guanosine | GRS | NT |
| Carnitine | 152 | AAD | GSH | GSH | AAD |
| 2-hydroxyglutarate | 2HG | TCA | Guanine | GUN | NT |
| Deoxyinosine | 2ND | NT | Phenyllactic acid | HFA | Others |
| 2PG | 2PG | Others | 1-Methyl-Histidine | HIC | AAD |
| Deoxyadenosine | 3D1 | NT | Hypoxanthine | HPA | NT |
| 3-Phospho-D-glycerate | 3PG | Carb. | 4-Hydroxyproline | HYP | AAD |
| Xanthosine | 4UO | NT | Indole-3-carboxylic acid | ICO | Others |
| GMP | 5GP | NT | 2-keto-isovalerate | KIV | AAD |
| GABA | ABU | AAD | Malate | LMR | TCA |
| acetylcarnitine | ACA | AAD | Methylcitrate | MCT | TCA |
| Adenine | AD0 | NT | 7-methylguanosine | MG7 | NT |
| Adenosine | ADN | NT | Malonic acid | MLA | Others |
| ADP | ADP | NT | Methylthioadenosine | MTA | AAD |
| alpha-ketoglutarate | AKG | TCA | NAD+ | NAD | NT |
| N6-Acetyl-L-lysine | ALY | AAD | NADH | NAI | NT |
| AMP | AMP | NT | Nicotinamide | NCA | Cofactors |
| N-Acetylorithine | AOR | AAD | NADP+ | NDP | NT |
| ATP | ATP | NT | N-acetylglutamate | NLG | AAD |
| N-acetylalanine | AYA | AAD | N-acetyl-glutamine | NLQ | AAD |
| betaine | BET | AAD | Inosine | NOS | NT |
| Biotin | BTN | Cofactors | Ornithine | ORN | AAD |
| C3-carnitine | C3C | AAD | Pantothenic acid | PAU | Cofactors |
| C4-carnitines | C4C | AAD | Phosphorylcholine | PC | AAD |
| C5-carnitines | C5C | AAD | Phosphoenolpyruvate | PEP | Others |
| CMP | C5P | NT | Phenylpyruvate | PPY | AAD |
| glycerophosphocholine | CH5 | AAD | Riboflavin | RBF | Cofactors |
| Cholic acid | CHD | Bile Acids | S-Adenosyl-L-methionine | SAM | Others |
| Choline | CHT | AAD | Saccharopine | SHR | AAD |
| Citrulline | CIT | AAD | Succinate | SIN | TCA |
| CoA | COA | Cofactors | Taurine | TAU | AAD |
| Creatine | CRN | AAD | cis-Aconitate | TRA | TCA |
| Cytidine | CTN | NT | Uracil | U0 | NT |
| Cystathionine | CTT | AAD | UMP | U5P | NT |
| dAMP | D5M | NT | UDP-GlcNAc | UD1 | NT |
| dCMP | DCM | NT | UDP | UDP | NT |
| Dimethylglycine | DMG | AAD | UDP-glucuronic acid | UGA | NT |
| Deoxycholic acid | DXC | Bile Acids | 2-aminoadipate | UN1 | AAD |
| FAD | FAD | Cofactors | UDP-hexose | UPG | NT |
| Flavone | FLN | Others | Uridine | URI | NT |
| Fumarate | FUM | TCA | Thiamine | VIB | Cofactors |
| Glycerol-3-phosphate | G3P | Carb. | N-Acetylputrescine | X5A | AAD |
| D-Glucose 6-phosphate | G6P | Carb. | Xanthine | XAN | NT |
| Deoxyguanosine | GNG | NT |  |  |  |

Class names: AAD: Amino Acid Derivatives; TCA: TCA Cycle; NT: Nucleotides; Carb.: Carbohydrates

**Supplementary Table S12.** Simulation details, including number of monomers (*N*_m_), box length (*L*_box_), and simulation time (*t*), for determining phase diagrams of PM and ATP mixtures at 298 K and 150 mM NaCl.

| PM:ATP ($\mu\text{M}:\text{mM}$) | $N_m$ (PM/ATP) | $L_{\text{box}}$ (nm) | $t$ ($\mu\text{s}$) |
| --- | --- | --- | --- |
| 3:1 | 100/33300 | 381 | 10 |
| 5:1 | 100/199000 | 321 | 10 |
| 10:1 | 100/10000 | 255 | 10 |
| 30:1 | 100/3339 | 177 | 10 |
| 100:1 | 100/1000 | 119 | 10 |
| 200:1 | 100/500 | 94 | 10 |
| 500:1 | 100/200 | 69 | 10 |
| 800:1 | 100/124 | 59 | 10 |
| 1000:1 | 100/100 | 55 | 10 |

### Supplementary Figures

**Supplementary Fig. S1.**
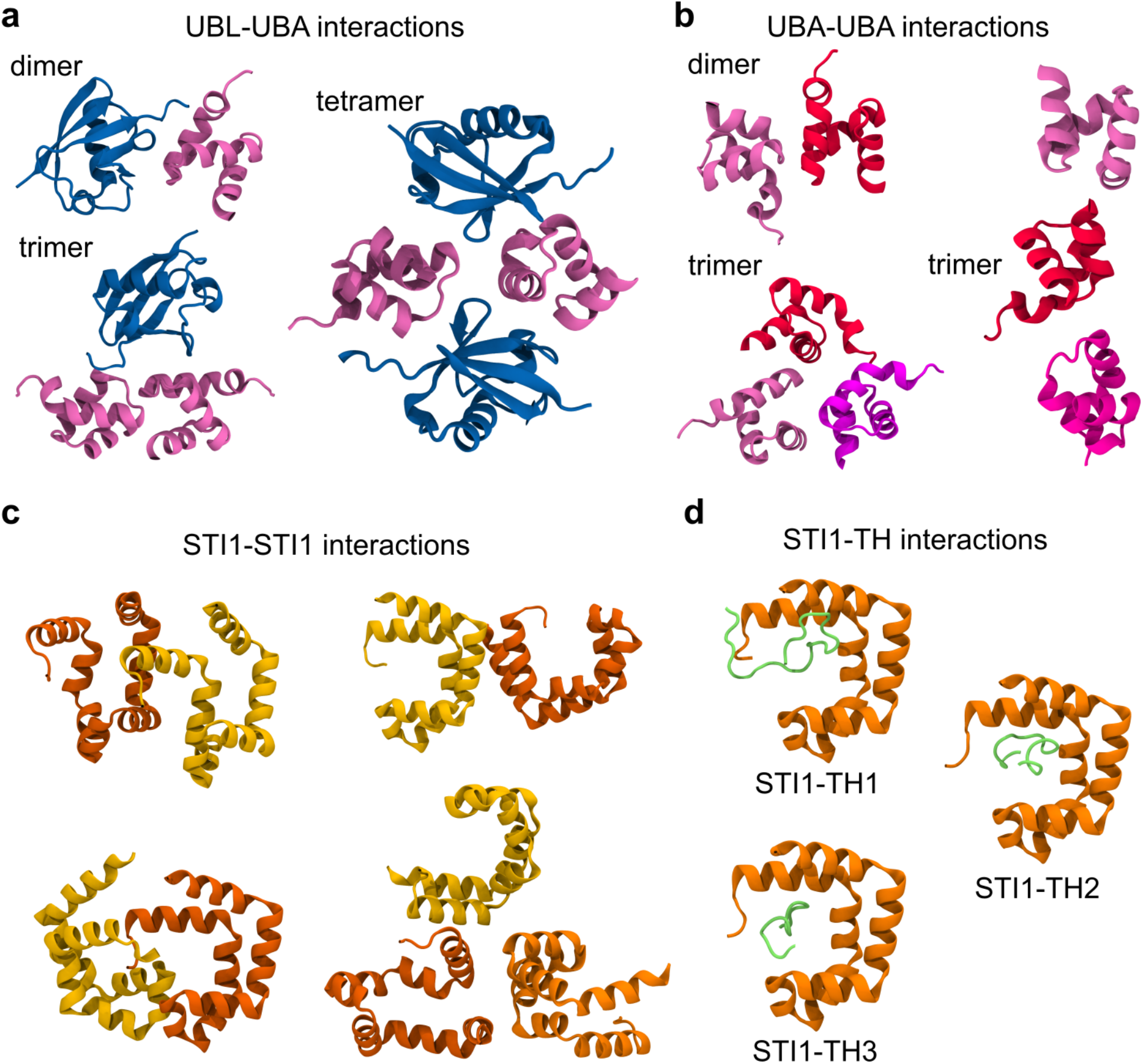
Representative snapshots of various intermolecular interactions between Dsk2 proteins, including UBL-UBA interactions (**a**), UBA-UBA interactions (**b**), STI1-STI1 interactions (**c**), and STI1-TH interactions (**d**).

**Supplementary Fig. S2.**
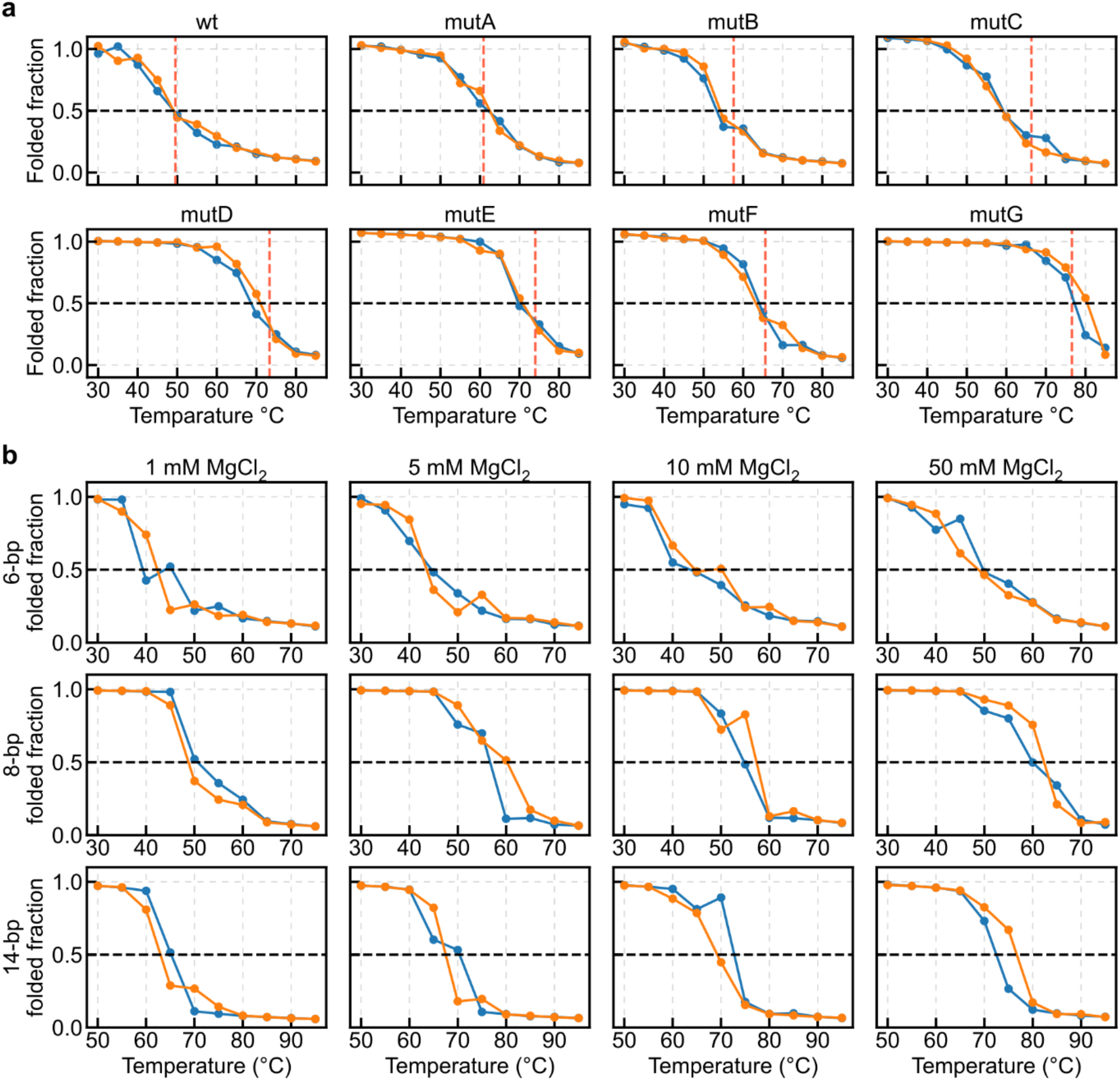
Determination of melting temperatures of RNA hairpins at 50 mM NaCl (**a)** and 6-bp, 8-bp, and 14-bp duplexes at 110 mM NaCl and 1, 5, 10, and 50 mM MgCl_2_ (**b**). Folded fraction was plotted against temperatures for two replicas. Dashed lines indicate the 50% folded state, which was used to estimate melting temperatures.

**Supplementary Fig. S3.**
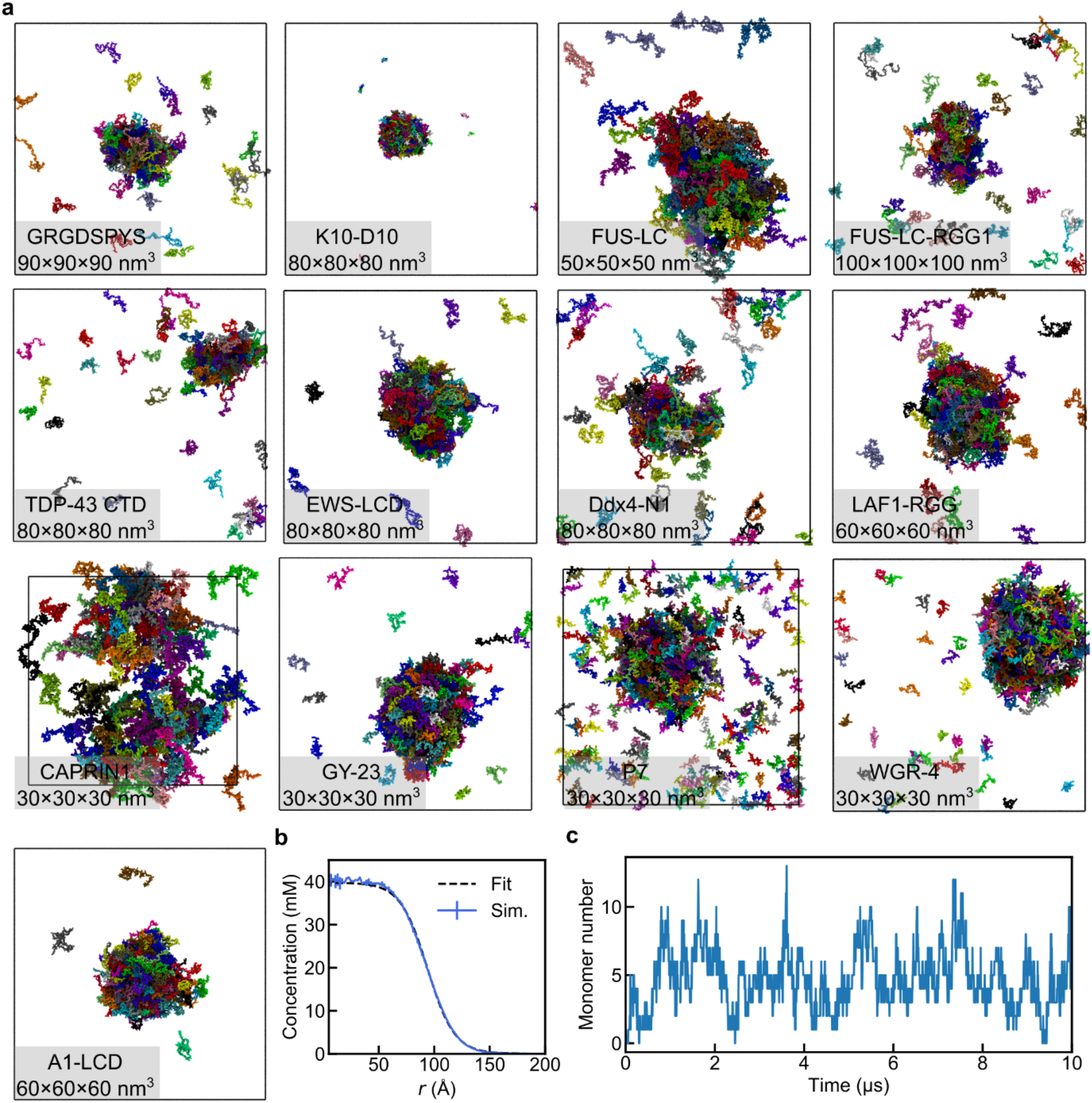
Direct phase coexistence simulations and determination of *C*_sat_. **a**, Representative final snapshots for the direct phase coexistence simulations of WT IDPs, with protein names and box dimensions indicated. The proteins are colored by chain identities. **b**, Density profile of the A1-LCD droplet (blue curve) and the corresponding fitted curve (Eq. 2; black dashed line). **c**, Number of A1-LCD proteins in the dilute phase as a function of the simulation time, showing strong convergence.

**Supplementary Fig. S4.**
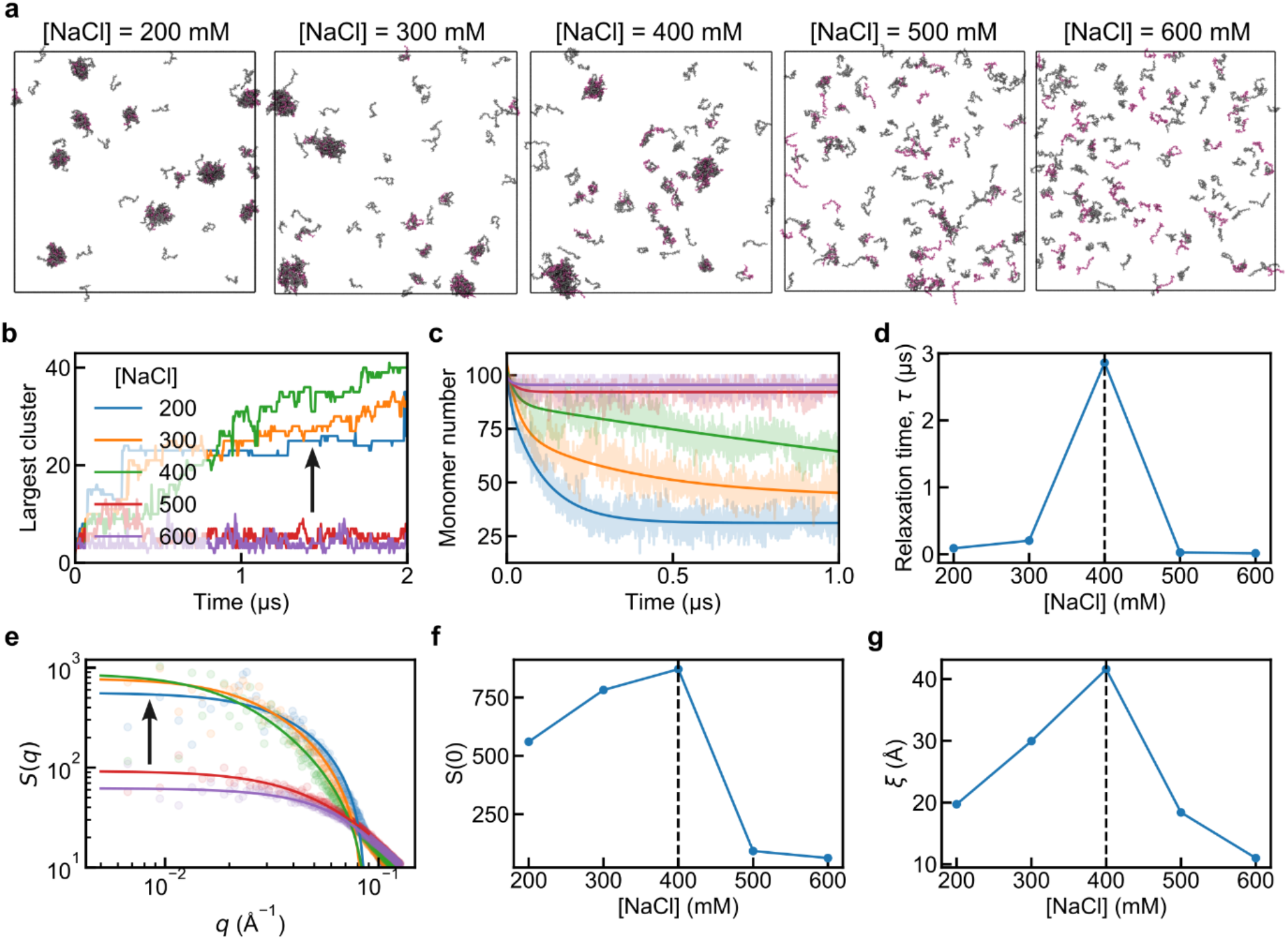
Determination of the phase boundaries of (PR)_30_ and rU_30_ mixtures by examining the cluster formation during modest 2-4 µs simulations. (**a**), largest cluster size (**b**), monomer costing (**c**) and its relaxation time (**d**), static structure factors (**e**) and derived density fluctuation amplitude (**f**) and correlation length (**g**). In panel **a**, peptides and RNAs are grey and mauve in color. In panels **c** and **e**, solid lines indicate the fitting curves. In panels **d, f**, and **e**, dashed lines indicate the transition points.

**Supplementary Fig. S5.**
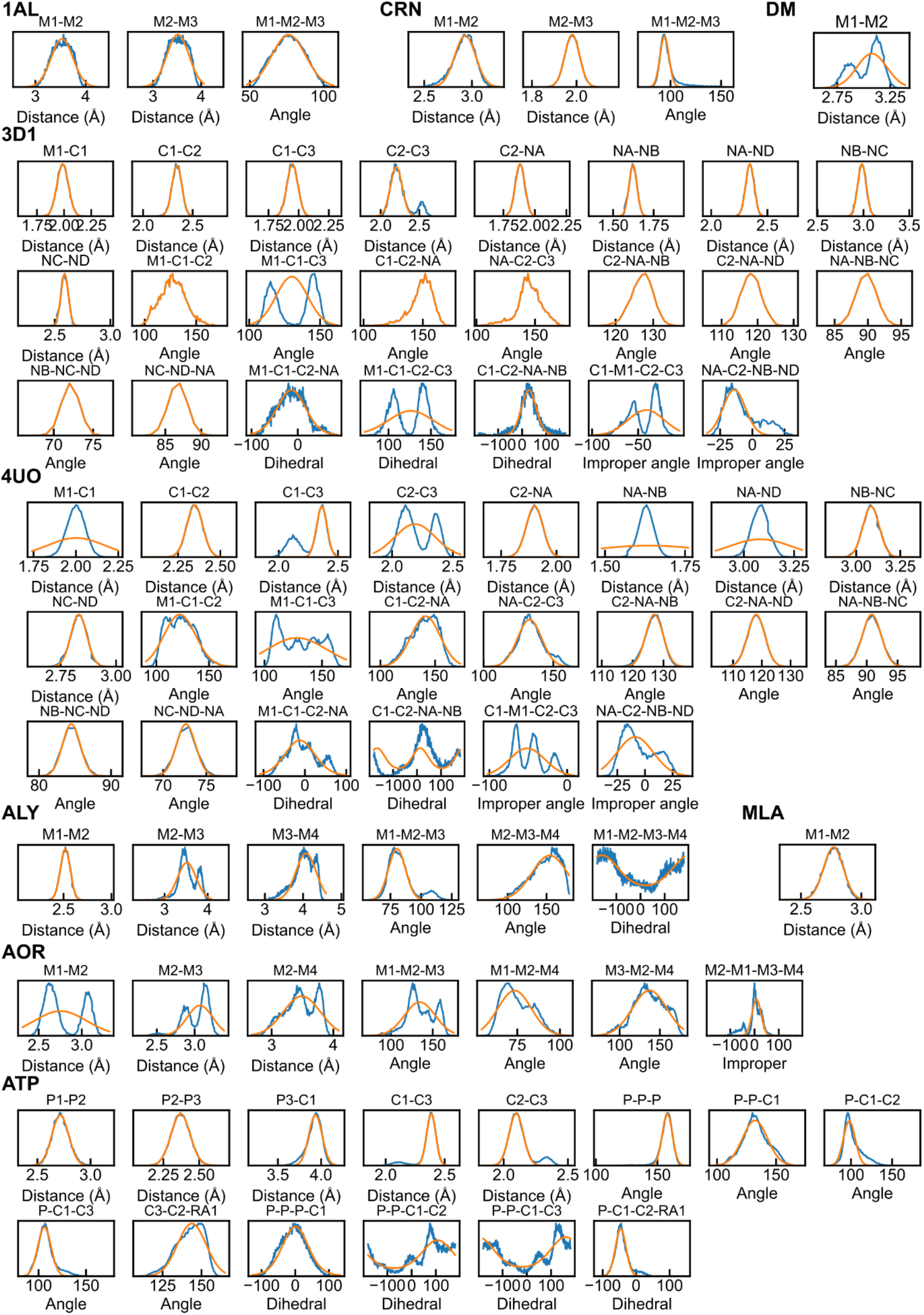
All-atom bonded distributions (blue) and the fitted results (orange) for 1AL, CRN, DM, 3D1, 4UO, ALY, MLA, AOR, and ATP.

**Supplementary Fig. S6.**
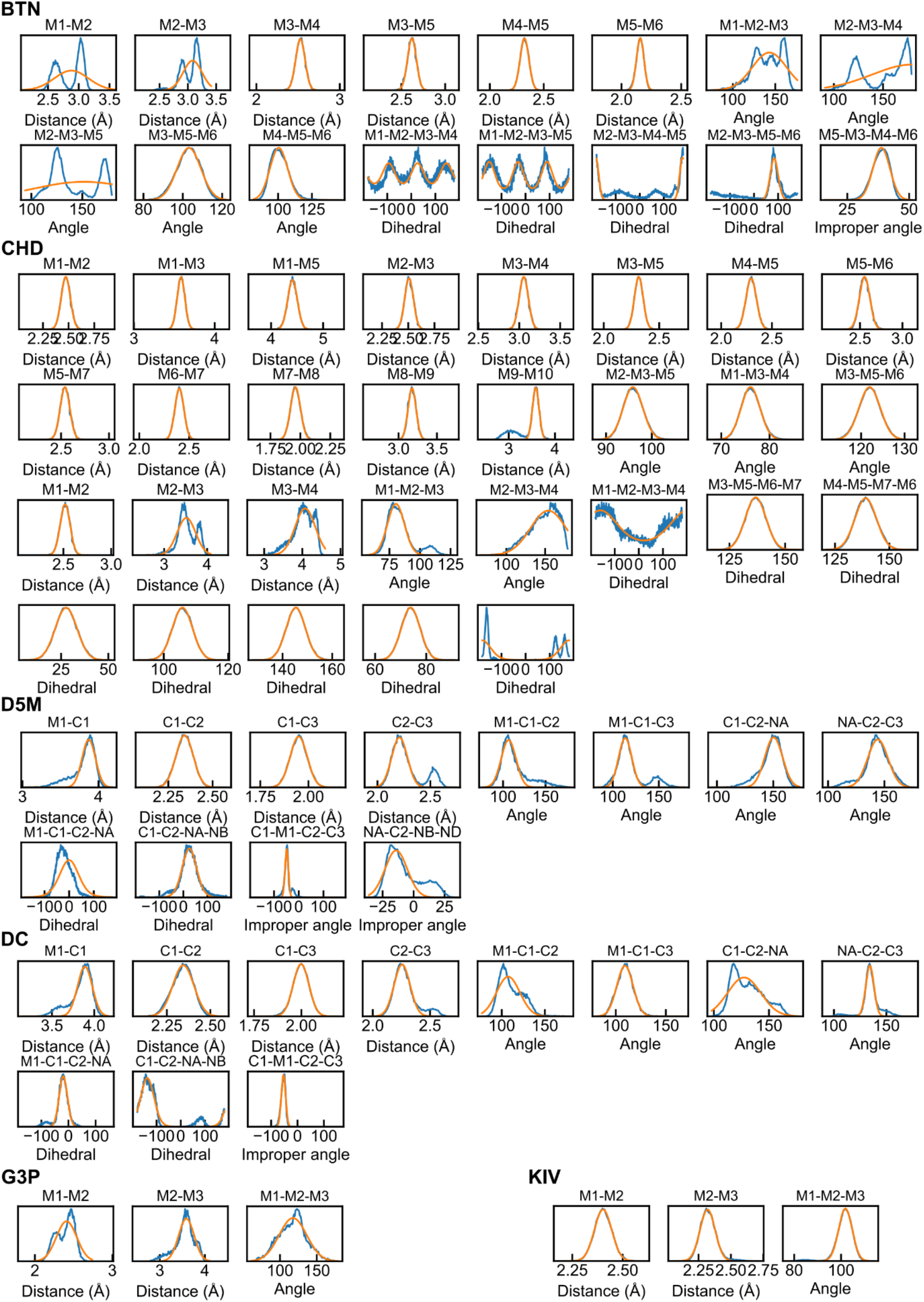
All-atom bonded distributions (blue) and the fitted results (orange) for BTN, CHD, D5M, DC, G3P, and KIV.

**Supplementary Fig. S7.**
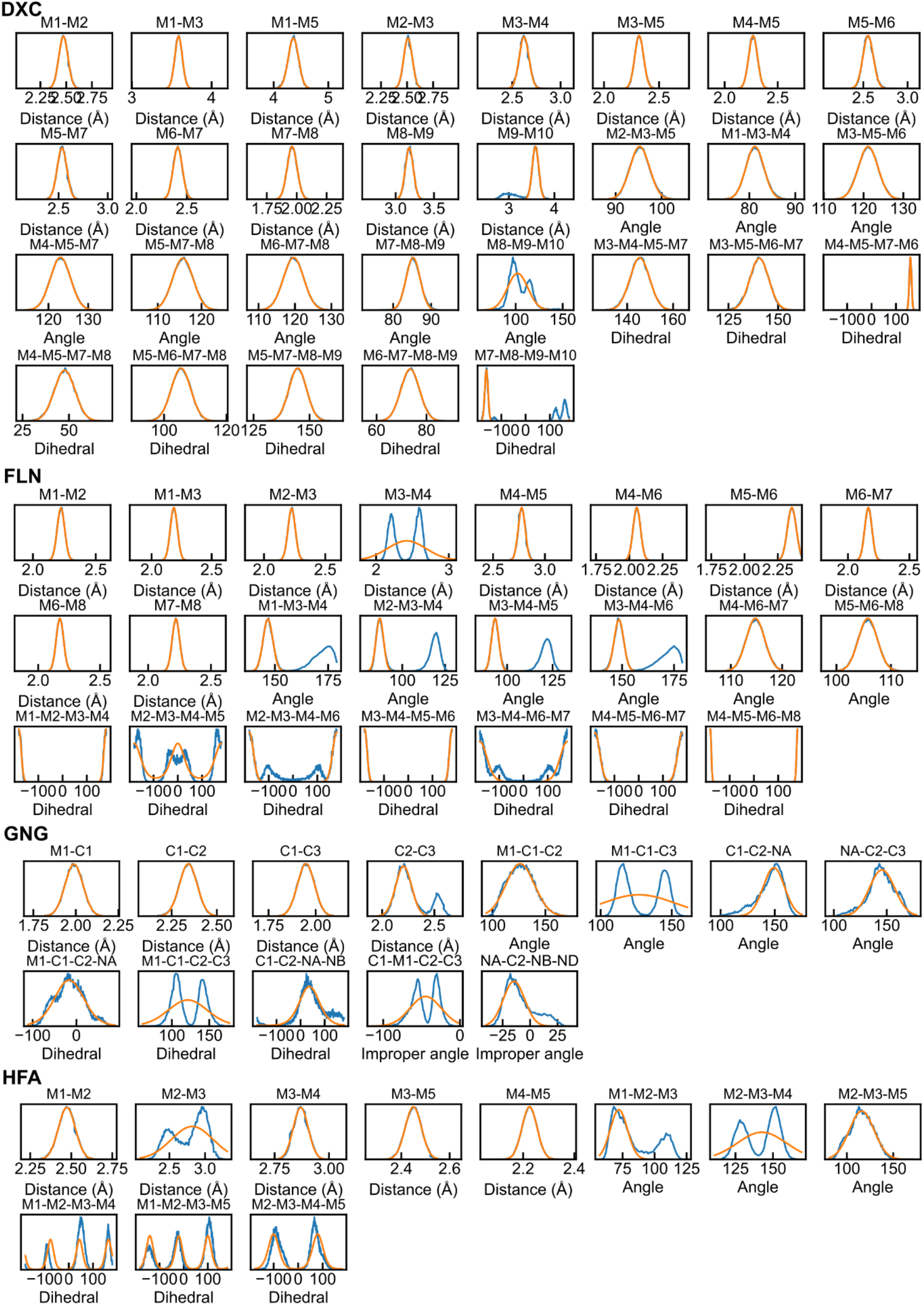
All-atom bonded distributions (blue) and the fitted results (orange) for DXC, FLN, GNG, and HFA.

**Supplementary Fig. S8.**
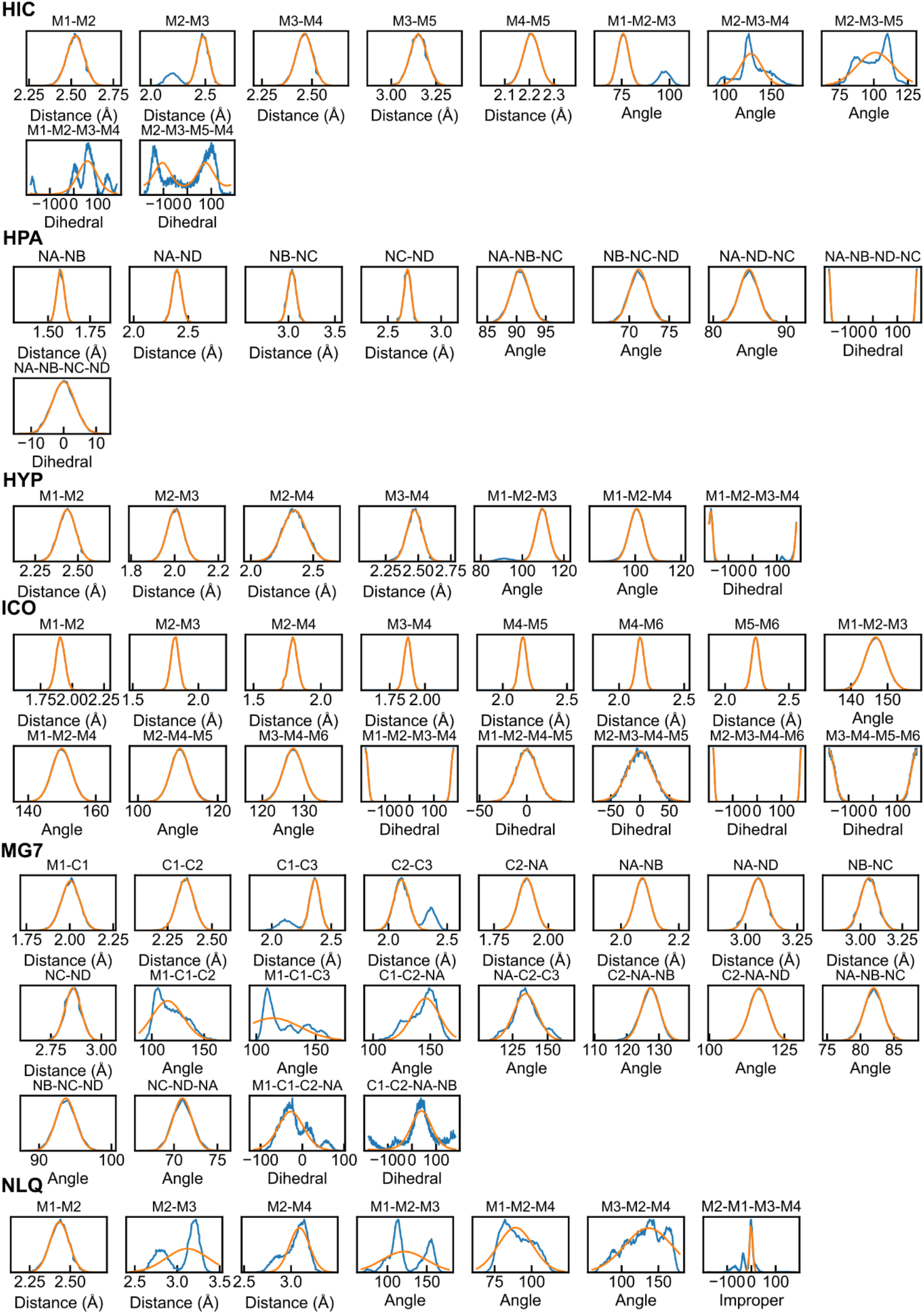
All-atom bonded distributions (blue) and the fitted results (orange) for HIC, HPA, HYP, ICO, MG7, and NLQ.

**Supplementary Fig. S9.**
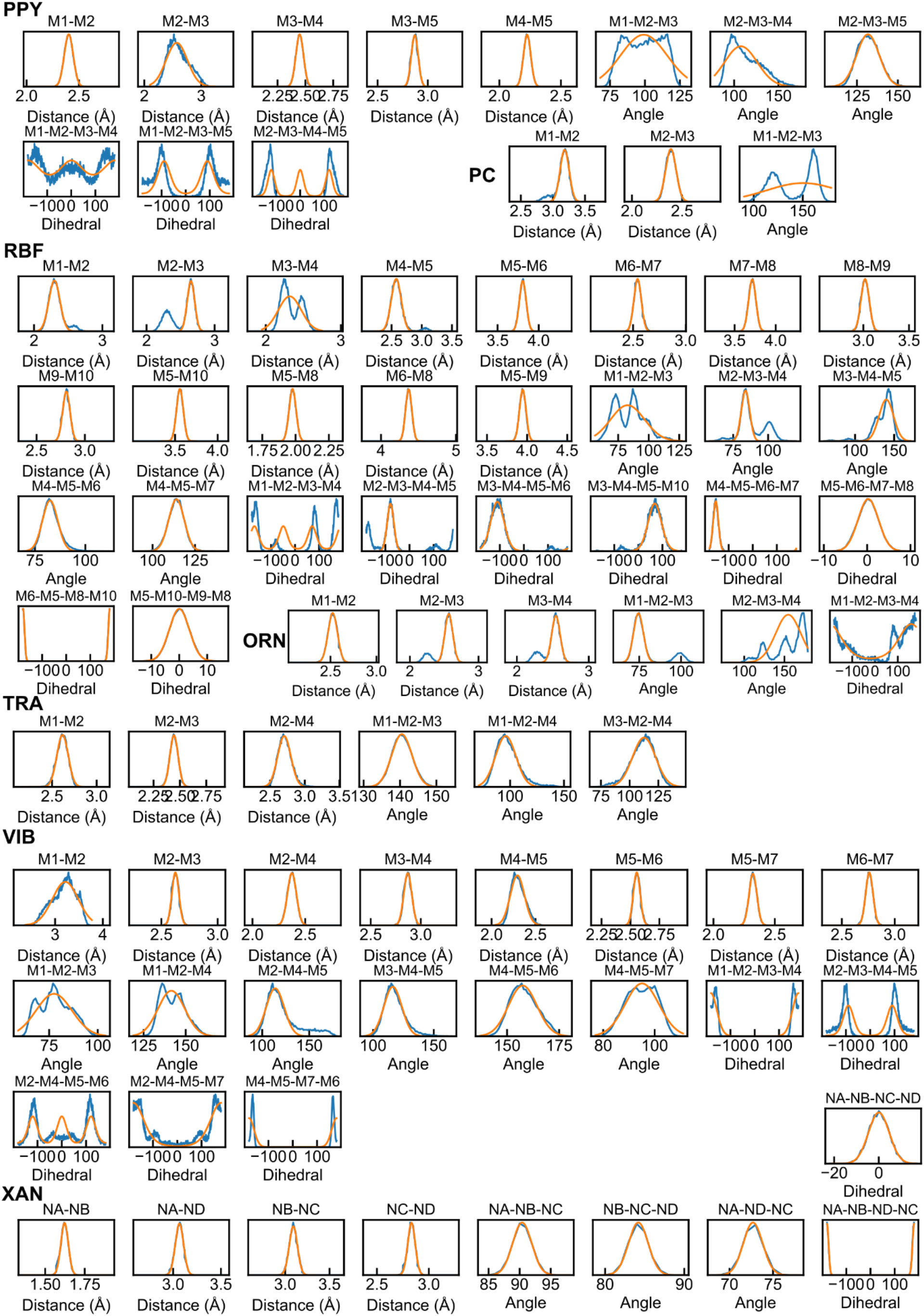
All-atom bonded distributions (blue) and the fitted results (orange) for PPY, PC, RBF, ORN, TRA, VIB, and XAN.

